# Identification of orphan ligand-receptor relationships using a cell-based CRISPRa enrichment screening platform

**DOI:** 10.1101/2022.06.22.497261

**Authors:** Dirk H. Siepe, Lukas T. Henneberg, Steven C. Wilson, Gaelen T. Hess, Michael C. Bassik, Kai Zinn, K. Christopher Garcia

## Abstract

Secreted proteins, which include cytokines, hormones and growth factors, are extracellular ligands that control key signaling pathways mediating cell-cell communication within and between tissues and organs. Many drugs target secreted ligands and their cell-surface receptors. Still, there are hundreds of secreted human proteins that either have no identified receptors (“orphans”) and are likely to act through cell surface receptors that have not yet been characterized. Discovery of secreted ligand-receptor interactions by high-throughput screening has been problematic, because the most commonly used high-throughput methods for protein-protein interaction (PPI) screening do not work well for extracellular interactions. Cell-based screening is a promising technology for definition of new ligand-receptor interactions, because multimerized ligands can enrich for cells expressing low affinity cell-surface receptors, and such methods do not require purification of receptor extracellular domains. Here, we present a proteo-genomic cell-based CRISPR activation (CRISPRa) enrichment screening platform employing customized pooled cell surface receptor sgRNA libraries in combination with a magnetic bead selection-based enrichment workflow for rapid, parallel ligand-receptor deorphanization. We curated 80 potentially high value orphan secreted proteins and ultimately screened 20 secreted ligands against two cell sgRNA libraries with targeted expression of all single-pass (TM1) or multi-pass (TM2+) receptors by CRISPRa. We identified previously unknown interactions in 12 of these screens, and validated several of them using surface plasmon resonance and/or cell binding. The newly deorphanized ligands include three receptor tyrosine phosphatase (RPTP) ligands and a chemokine like protein that binds to killer cell inhibitory receptors (KIR’s). These new interactions provide a resource for future investigations of interactions between the human secreted and membrane proteomes.

## INTRODUCTION

The human proteome can be envisioned as an array of nodes grouped into local communities, where each node represents one protein and each local community represents a protein complex or network (Budayeva and Kirkpatrick, 2020; Huttlin et al., 2017). These communities determine physiological function and subcellular localization. Many communities include secreted protein ligands, their cell-surface receptors, and signaling molecules that bind to the receptors. The human secretome on its own constitutes approximately 15% of all human genes and encodes more than 4000 different proteins (Uhlén et al., 2019) with a wide range of tissue expression (Figure 1B). Most of the new drugs developed in recent years target secreted proteins and their receptors, and new therapeutic targets are likely to emerge from screens to identify ligand-receptor interactions (Clark et al., 2003; Stastna and Van Eyk, 2012).

**Figure 1.**
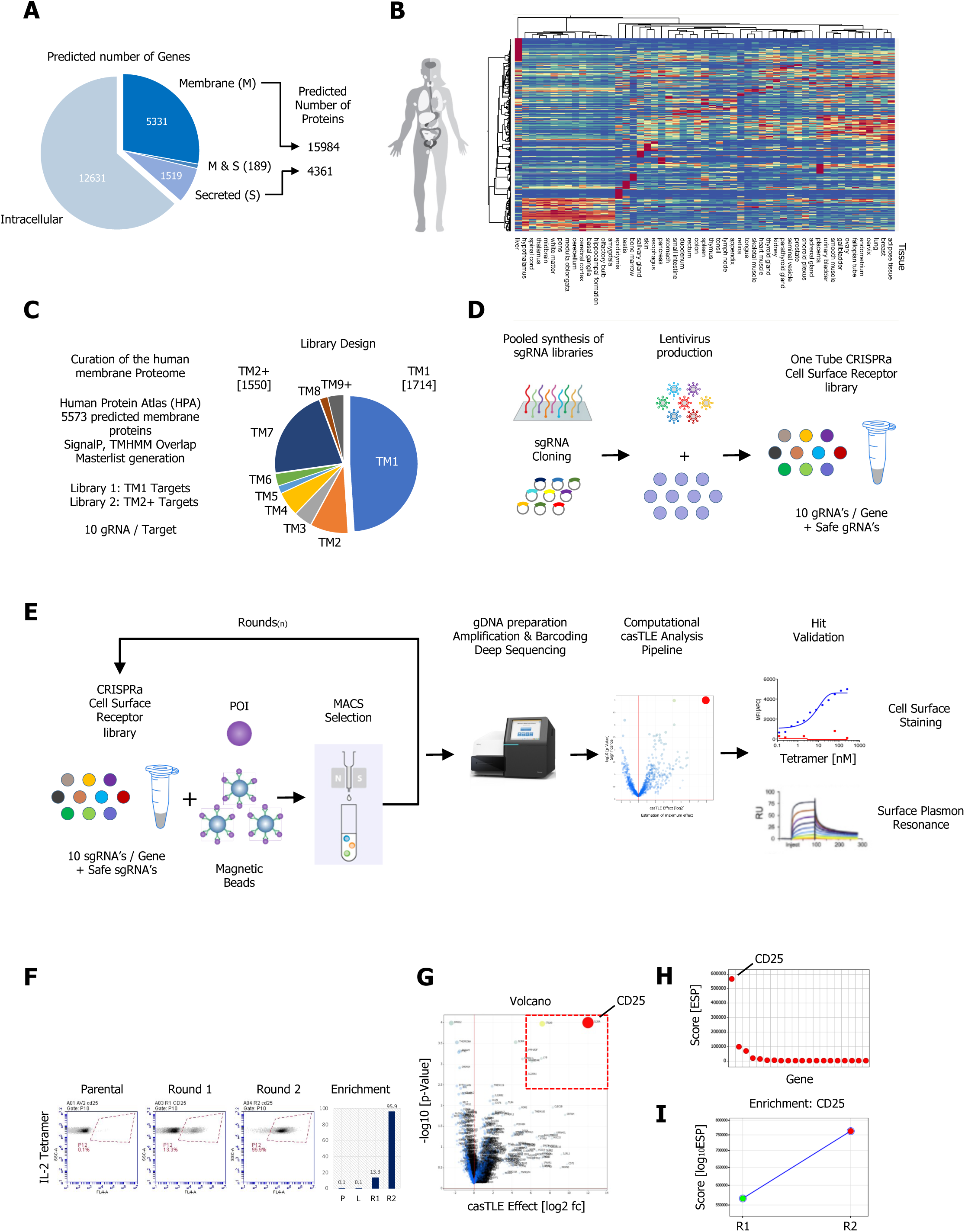
A CRISPR activating enrichment screening platform. Curation of the human membrane proteome, cell surface Library design, validation and benchmark screen. (A) Human membrane and secreted proteome; left panel: predicted number of intracellular, membrane (M) and secreted (S) genes with a total number of approximately 5520 human protein-coding genes predicted to encode ∼15984 membrane-spanning proteins including mapped, alternative splice variants and isoforms. (B) Secreted proteome visualized by 2-way hierarchical clustering of normalized mRNA expression data from normal tissue (C) Human membrane proteome curation and workflow of the cell surface library design. (D) Pooled, customized and target-specific TM1 and TM2+ sgRNA libraries (10 sgRNA/target) were designed, cloned and lentivirally infected into K562-CRISPRa cells at low MOI. (E) Schematic overview of the CRISPRa enrichment screening platform; A Protein of interest (POI) is complexed with magnetic beads and screened against customized CRISPRa cell surface receptor library, followed by consecutive rounds of MACS positive selection. In the final step, genomic DNA is extracted from the selected, target enriched library round(s), barcoded, subjected to deep sequencing and analyzed using the casTLE statistical framework to identify potential hits. CRISPRA hits are then subjected to various orthogonal validation methods. (F-I) Benchmark CRISPRa enrichment screen using human IL-2, performing 2 consecutive rounds of magnetic bead selection followed by gDNA extraction, barcoding and deep sequencing. (F) Enrichment over 2 rounds of consecutive magnetic bead selection by tetramer staining with human IL-2 post selection (Parental, Round 1 and Round 2). (G) Visualization of the deep sequencing analysis. Results are visualized by x/y scatter plot: casTLE-Score (log2); pValue (-log10), size of the hit represents the casTLE-Effect + casTLE-Score. (H) Candidate hits of the final round of enrichment visualized by a x/y ranked plot using a combined ESP score (log10). (I) Trajectories of highest ranking candidates are plotted over the consecutive rounds of enrichment rounds, size of the bubble represents the p-Value (-log10).

Mapping of interactions that occur at the cell surface has significantly lagged behind that of intracellular interactions, because the most widely used high-throughput protein-protein interaction (PPi) screening methods, including affinity purification/mass spectrometry (AP/MS), yeast two-hybrid screening (Y2H), and phage display, are not well suited to analysis of extracellular domain (ECD) interactions (Havugimana et al., 2012; Huttlin et al., 2015; Krogan et al., 2006; Martinez-Martin, 2017). ECD interactions are often of low affinity, with K_D_s in the micromolar range, and can have fast dissociation rates, rendering them difficult to detect since they may not produce stable complexes (Honig and Shapiro, 2020). As a consequence, ECD interactions are generally underrepresented in screens that rely on the formation of such complexes (Braun et al., 2009; Martinez-Martin et al., 2019; Özkan et al., 2013c; Söllner and Wright, 2009; Wojtowicz et al., 2020). In addition, many putative ECD interactions reported by AP/MS and Y2H protein interaction databases have the tendency to be false positives. AP/MS produces false positives for cell-surface proteins due to incomplete solubilization of membranes, leading to identification of indirect interactions. Y2H examines interactions inside the cell, but most ECDs have disulfide bonds and glycosylation sites. To acquire these modifications and fold correctly, cell-surface and secreted proteins must move through the secretory pathway. Because of this, ECD interactions detected by Y2H are often false positives due to domain misfolding. Similar issues apply to phage display and to microarrays in which mRNAs are translated on a chip. Thus, while these high-throughput methods can identify interactions with the cytoplasmic domains of receptors, they usually fail to find genuine ECD interactions.

Successful high-throughput screens to detect weak ECD interactions in vitro have taken advantage of avidity effects by expressing ECDs as fusions with multimerization domains. In such binary interaction screens, one protein (the bait) is applied to a surface, and the other (the prey) is in solution. Prey binding to the bait is assessed using colorimetric or fluorescent detection. These methods include AVEXIS, apECIA, alpha-Screen and BPIA, which are carried out using ELISA plates, chips, or beads (Braun et al., 2009; Bushell et al., 2008; Li et al., 2017; Martinez-Martin, 2017; Taouji et al., 2009). However, in vitro screens have limitations. They require robotic high-throughput instrumentation and are time-consuming and expensive to carry out on a large scale, since they require synthesis of ECD coding regions and expression of individual bait and prey proteins. In addition, in vitro screens cannot usually assess binding to ECDs of receptors that span the membrane multiple times, because such ECDs are often composed of noncontiguous loops and cannot be easily expressed in a soluble form. Furthermore, in vitro binary interaction mapping technologies lack the natural spatial context of the cell membrane. They may also fail in cases where cofactors and/or post-translational modifications are required for binding.

To address these issues, several groups have developed cell-based screens for receptor-ligand interactions that take advantage of CRISPR technology (Cong et al., 2013; Jinek et al., 2013; Mali et al., 2013). In CRISPR activation (CRISPRa) screens such as the one described here, induce gene expression by targeting transcriptional activators to their control elements using sgRNAs (Chong et al., 2018; Kampmann, 2018; Morgens et al., 2016; Tanenbaum et al., 2014a). Utilizing CRISPRa pooled sgRNA libraries eliminates the need to create expensive collection of synthetic genes, and in addition allows a forward positive screening workflow which enables a higher dynamic range compared to loss of function screens (Doench, 2018). Libraries of cells, each with an sgRNA targeting one receptor, can be easily stored and screened for binding to soluble ligands.

Here we describe a CRISPRa enrichment workflow that employs customized, pooled cell surface receptor sgRNA libraries in combination with magnetic bead-based selection (MACS) to enrich for receptor-expressing cells. This approach allows cost-efficient parallel screening with multiple ligands. We created two cell libraries, comprising all single-transmembrane (TM1) and multi-pass transmembrane (TM2+) receptors, and screened them with a collection of secreted ligands. To define a set of high-priority ligands, we first curated the human secreted proteome and selected and expressed 20 at levels sufficient for screening the TM1 and TM2+ libraries. We identified new receptor candidates in more than half of these screens. These were validated using surface plasmon resonance (SPR) and/or cell binding. These studies define new receptors for several secreted ligands that function in the immune and nervous systems.

## RESULTS

### A CRISPR activating enrichment screening platform

CRISPRa mediated activation of transcription using the sunCas9 system is a precise and scalable method for inducing expression of endogenous genes across a high dynamic range (Gilbert et al., 2014a). This system uses a dead Cas9 (dCas9) variant fused to a SunTag, a multicopy epitope tag that recruits the VP64 transcriptional activator via binding to a cytoplasmic scFV-nanobody-VP64 fusion protein. sgRNAs guide this complex to the enhancer region of the gene of interest and facilitate target-specific gene activation and expression (Tanenbaum et al., 2014b).

To evaluate the performance and feasibility of CRISPRa mediated transcriptional activation of cell surface proteins for a receptor/ligand interaction discovery platform, we first selected 10 well-characterized cell surface receptors with varying mRNA expression levels ranging from not detected to highly expressed in K562 human myeloid leukemia cells (Figure S1A) (Thul et al., 2017; Uhlén et al., 2019). We then generated a pooled lentiviral mini-library of 10 sgRNAs per enhancer (100 sgRNA elements), matched with 100 control sgRNAs derived from scrambled sequences (Gilbert et al., 2014a) and transduced K562 cells stably expressing the sunCas9 system (Figure S1A). Each library plasmid contained a single sgRNA targeting one of the 10 genes, a GFP fluorescent marker and a puromycin resistance marker. The library transduced cells were puromycin selected for 5 days to obtain >90% GFP positive cells. The expression levels of the 10 cell surface receptors were then evaluated by cell surface staining using APC (Allophycocyanin) labelled antibodies against the respective targets (CD122 was used as a control). All 10 selected targets showed elevated cell surface expression in comparison to non-transduced K562 sunCas9 cells or a control receptor (CD122) (Figure S1B).

We then used human interleukin 2 (IL-2), which has a high affinity receptor subunit termed CD25, to validate our screening workflow (Figure 1E) in two parallel screens, simulating two library sizes by diluting the 10 target (100 sgRNA) K562 sunCas9 mini-library by 1:20 and 1:200 with non-transduced K562 sunCas9 cells, corresponding to final library sizes that would correspond to screening of 200 and 2000 targets, respectively (Figure S1C, D). Both library pools were incubated with magnetic streptavidin microbeads complexed with biotinylated IL-2, and IL-2 binding cells were isolated in a positive selection workflow by MACS, using Miltenyi LS-MACS columns (Figure S1E). After labelling, washing and elution, positively selected cells were expanded and stained with IL-2 tetramers (Figure S1D). Genomic DNA was extracted from both consecutive rounds of selection as well as the K562 sunCas9 ML library itself, followed by barcoding and deep sequencing for both libraries.

Deep sequencing data for each round of selection was analyzed and hits were identified using the robust casTLE statistical framework (Morgens et al., 2016b). Briefly, casTLE compares each set of gene-targeting guides to the negative controls, using both safe-targeting and non-targeting controls and selecting the most likely maximum effect size (casTLE-Effect). A *p*-value is then generated, representing the significance of this maximum effect by permuting the results. We calculated casTLE metrics for each round of selection in comparison to the naïve library. In addition, we used the casTLE metrics to plot trajectories for each hit for the consecutive rounds of selections, which allows for direct evaluation of sgRNA enrichment throughout the selection workflow and easy elimination of false positives. Using casTLE, both IL-2 CRISPRa screens successfully identified interleukin 2 (IL-2) receptor alpha (IL2RA; CD25) as the top hit with the highest confidence (casTLE Score), casTLE Effects and significance (*p*-value) (Figure S1E). Side by side comparison of enrichment scores for both rounds of selections from both libraries was plotted as bar graphs (Figure S1F).

### Customized, pooled CRISPRa cell surface receptor library design

Having established the screening workflow (Figure 1E), we sought to leverage the power and efficiency of customized, pooled CRISPRa cell surface libraries to perform targeted screens with secreted ligands. We first compiled a comprehensive list of cell surface receptors by carefully curating the human membrane proteome (Figure 1A, C). We choose a targeted cell surface library approach instead of a genome-wide approach, because it allowed for a smaller library size, resulting in a better signal to noise ratio (SNR) and avoiding unwanted transcriptional upregulation of non-membrane proteins. We utilized several databases including HUGO, UniProt, the Human Protein Atlas and bioinformatic tools (SignalP, TMHMM) to compile two cell surface target lists covering both single transmembrane (TM1) and multi-span transmembrane (TM2+) cell surface proteins (Figure 1C). For the CRISPRa mediated transcriptional activation of cell surface proteins, we synthesized and cloned two pooled sgRNA libraries, a TM1 and a TM2+ library, each with 10 sgRNAs per target (Gilbert et al., 2014b). Both libraries include matched controls targeting genomic locations without annotated function (Figures 1C, D). K562 cells stably expressing the sunCas9 system were infected with both libraries (TM1; TM2+) at low, medium and high multiplicity of infection (MOI), then selected with puromycin until the cell population was at least 90% GFP-positive, indicating the presence of lentivirus. Cells were recovered and expanded, and representative aliquots were saved as naïve library stocks in liquid nitrogen with at least 1000x cell number coverage per sgRNA to maintain maximum library complexity. Sufficient sgRNA representation of the naïve library was confirmed by deep sequencing after selection and showed the highest coverage and diversity at low MOI with at least 91% of reads with at least one reported alignment (R=0.97) for both libraries (Figure S1G, H). Library information including sgRNA target IDs and sequences for both libraries (TM1; TM2+) can be found in Data S3.

### CRISPRa benchmark screen using human interleukin 2 (IL-2)

After library cloning and validation, we benchmarked the sensitivity and robustness of our screening platform with a proof of concept screen using human interleukin 2 (IL-2), following the protocols used for the mini-library (Figure S1). We successfully recovered CD25 (IL2RA) as the top hit after two rounds of enrichment, deep sequencing and analysis following the outlined workflow (Figure 1E). Initially, the naïve TM1 library showed no positive IL-2 binding by tetramer staining (Figure 1F). Library enrichment was monitored by IL-2 tetramer staining throughout the selection workflow and only after one round of positive selection we observed a significant enrichment of IL-2 selected cells from 0.1% to 13.3% IL-2 tetramer positive cells (Figure 1F). After expanding the cells from the first round and subjecting them to a second round of CRISPRa enrichment screening we observed a further robust increase of IL-2 tetramer positive from 13.3 to 95% IL-2 tetramer positive cells (Figure 1F).

After each consecutive round of selection, enriched cells were expanded and genomic DNA was extracted, followed by barcoding and deep sequencing. Genomic DNA from the K562 sunCas9 TM1 naïve library itself served as the baseline. Following deep sequencing, data from both rounds of consecutive IL-2 selections was analyzed and visualized using the casTLE statistical framework. CD25 was identified as the top hit with the highest confidence (casTLE Score), *p*-value (significance), and casTLE Effect (Figure 1G, H). Furthermore, we used the casTLE metrics from each round to plot trajectories of CD25, which allows for a direct evaluation of sgRNA enrichment throughout the selection workflow and shows a positive trajectory for CD25 in the selection workflow (Figure 1I), validating the sensitivity and robustness of our screening pipeline.

### Selection and production of secreted proteins for CRISPRa screening

We first generated a secreted proteome master list from several databases, including HUGO, UniProt, the Human Protein Atlas (Uhlén et al., 2019) and bioinformatic tools (SignalP, TMHMM) to identify potential high priority secreted proteins for our screening workflow. After curation of the human secreted proteome (Figure 1A, B), approximately 60% of the ∼1600 genes were classified as encoding enzymes (mostly proteases), enzyme inhibitors, serum proteins or components of saliva, tears, or other fluids (these include carrier proteins), structural, extracellular matrix proteins, antimicrobial, complement factors, coagulation factors, lectins, or unknown. The remaining ∼40% of genes were identified as likely to encode secreted ligands acting through cell surface receptors and further examined through literature searches. We classified products of 419 genes as ligands with known receptors that can adequately account for their biology. Finally, we identified 206 gene products either as “orphans” with no identified receptor or as ligands that are likely to have additional, as yet unidentified receptors in addition to those that have been described. From these 206, we ultimately selected a total of 80 high priority targets (one per gene; we did not consider isoforms generated through alternative splicing) with a broad coverage of molecular function, tissue expression, domain architecture, and disease association (DisGeNet) (Figure S2A, B). Coding sequences for these 80 secreted proteins of interest were synthesized, subcloned into an Avi-6xHIS expression plasmid, expressed in Expi293F cells, purified with Ni-NTA resin, then biotinylated in vitro and further purified by size-exclusion chromatography (SEC) (Figure S8A).

### CRISRPa enrichment screens reveal new secreted ligand-receptor interactions

We obtained sufficiently high expression levels for 20 of the 80 high priority targets (Figure 2A, S8A) to allow screening using our enrichment workflow (Figure 1E). Names and mRNA expression patterns in normal tissue for these 20 ligands are shown in Figures 2A. Each of the 20 was used to screen the TM1 and TM2+ libraries with up to 3 consecutive rounds of selection, followed by deep sequencing and statistical analysis using the casTLE framework. Screening results of the final round of enrichment for each of the 20 secreted proteins were filtered using the following cut-offs: casTLE-Effect > 2, casTLE-Score > 2, pValue < 0.05. To predict high-confidence interaction pairs from each dataset, a custom score was computed for each potential interaction pair by combining three metrics into one ESP score: (casTLE-Effect + casTLE Score)/p-Value. To integrate data analysis and visualization, we used the combined ESP score to rank sort interaction pairs for every screen.

**Figure 2.**
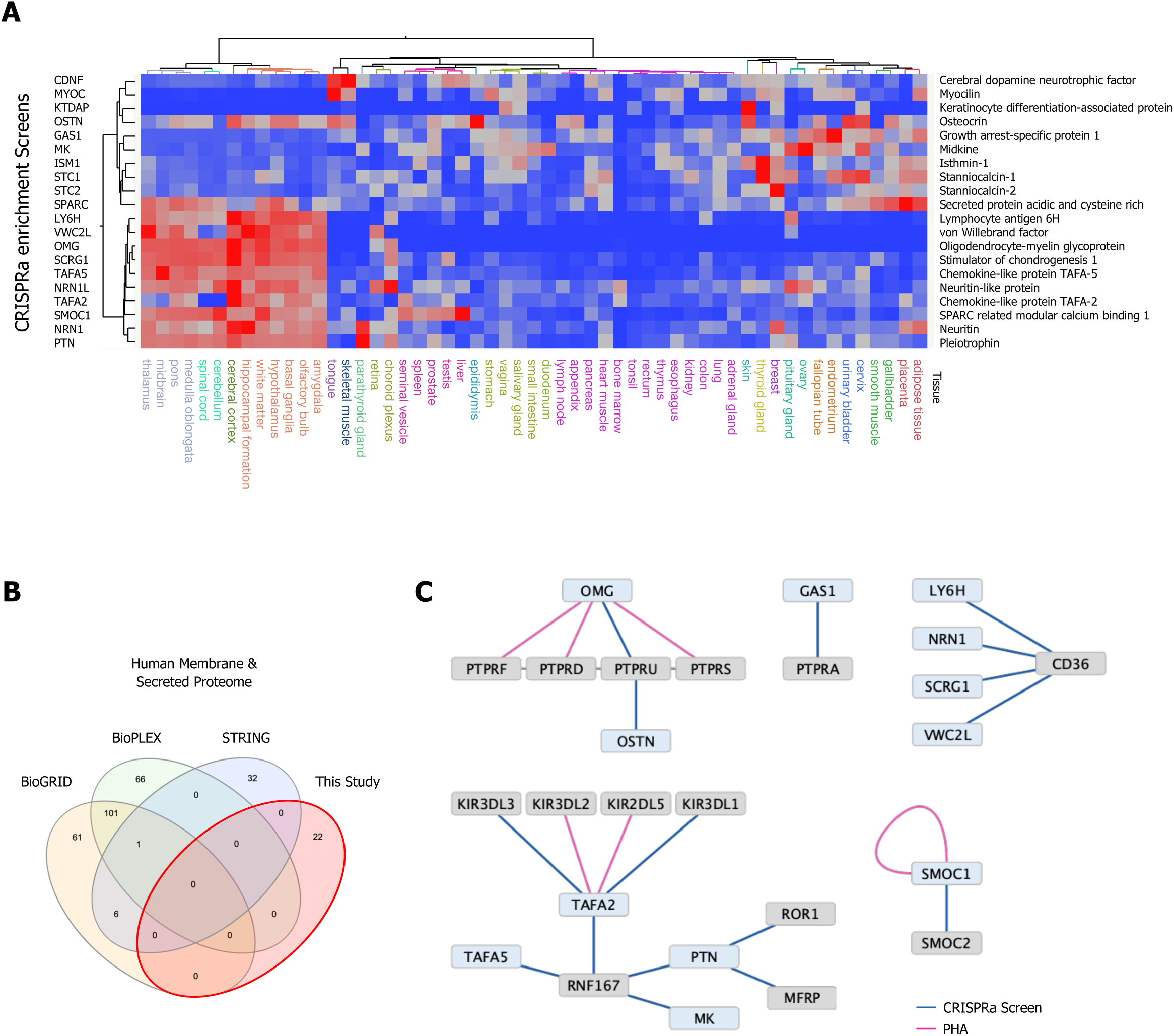
Selection and protein production of high priority secreted ligands and overview of screening results. (A) Curation of the secreted human proteome (Figure 1B) and high value target selection. Out of a total of ∼1600 predicted secreted proteins, 80 high value orphan secreted proteins were selected, synthesized, cloned and expressed in Expi293 cells. 20 secreted ligands with a wide range of tissue expression (A) were screened using our CRISPRa enrichment workflow (Figure 1E). (B) Venn diagram visualizing the overlap between physical interactions between secreted and membrane proteins presented in this study and public interaction databases (BioGRID, BioPLEX and STRING). (C) CRISPRa enrichment screening interactions represented as a protein interaction network, nodes represent CRISPRa query secreted ligands (blue) and candidate hits (grey). Edges represent the interactions between nodes. The visualized network shows 22 interactions between secreted and membrane proteins; 18 new interactions from 8 screens in the TM1 and 4 interactions from 4 screens in the TM2+ library between secreted and membrane proteins. Interactions (edges) resulting from CRISPRa enrichment screens are represented in dark blue, interactions resulting from phylogenetic homology analysis (PHA) are visualized in purple.

We selected a subset of CRISPRa enrichment screens with high ranking predicted ESP scores for potential interaction pairs for further validation using orthogonal methods, including surface plasmon resonance (SPR) and cell surface staining (CSS). Cell surface staining was utilized as an orthogonal validation method to show PPIs in a cellular context using a fluorescent-tetramerization-based approach by flow cytometry for high sensitivity detection of putative PPI’s on the cell surface. In total, we tested 22 candidate PPIs between the secreted and membrane proteome by SPR and/or CSS from 12 screens with PPIs in both the TM1 and the TM2+ library (Table 1). These validation data are shown for selected PPIs in Figures 3-6 and S6-S7.

**Figure 3.**
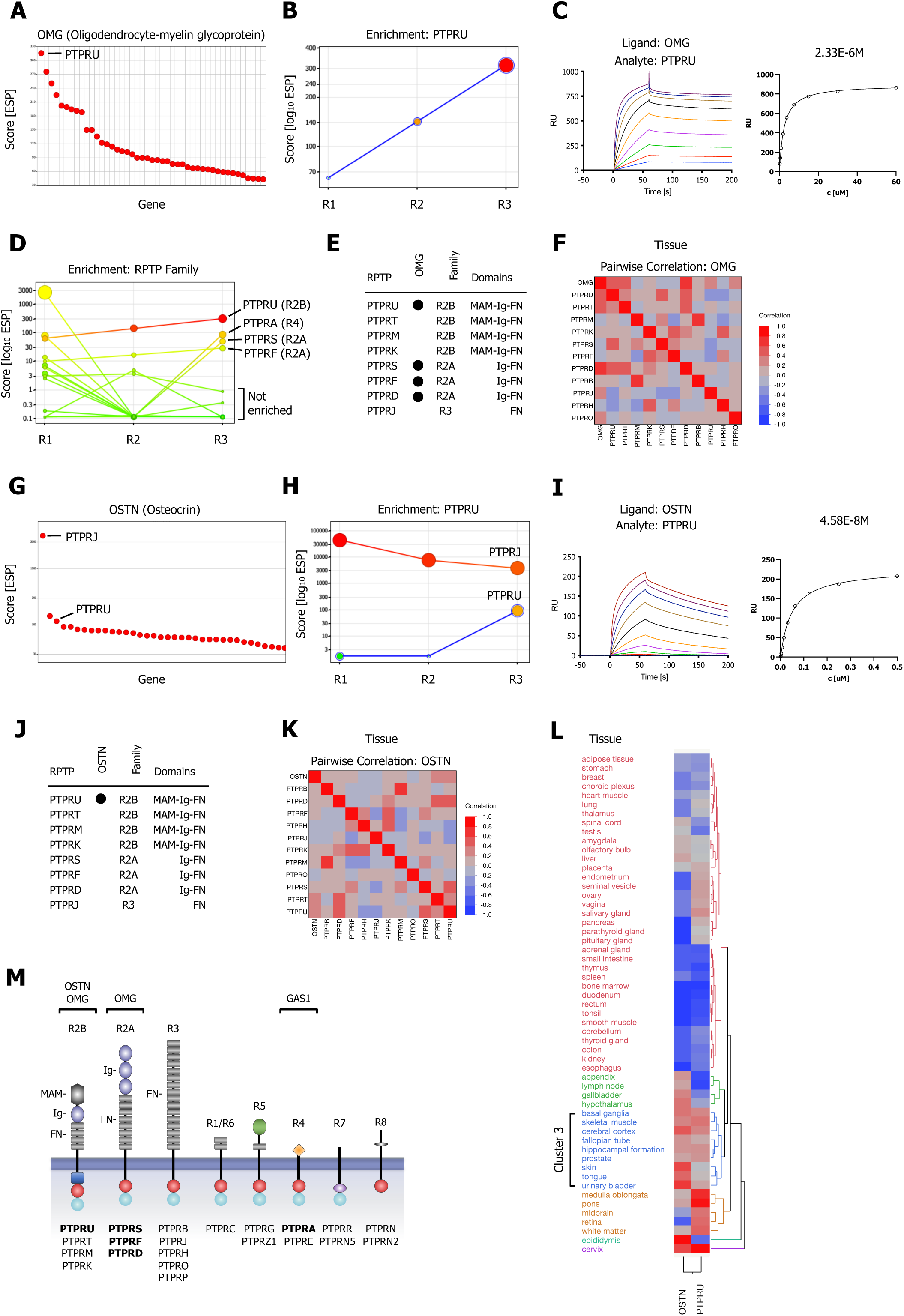
CRISPRa screening results and SPR Validation of Type R2A and Type R2B PTPRs for OMG and OSTN. (A) Ranked x/y scatter plot of Round 3 of the OMG screen. (B) Depicts the trajectory of the highest ranking candidate, PTPRU, plotting ESP scores for all 3 consecutive rounds of selections in a x/y enrichment plot. The size of the bubble represents the *p*-value (-log10). (C) SPR sensorgram and steady-state curve for human PTPRU-ECD (analyte) binding to human OMG (ligand). (D) Shows enrichment scores for additional members of the RPTP family members found in the OMG screen. (E) Summary of SPR results testing binding of R2A, R2B and R3 RPTP subfamily members (Figure S3A). (F) Multivariate heatmaps for OMG calculated from normal tissue mRNA expression correlations. (G) OSTN screen ranked x/y scatter plot of Round 3 ESP scores. (H) Trajectory of high ranking candidates PTPRJ and PTPRU by plotting ESP scores for all 3 consecutive rounds of selections in a x/y enrichment plot; size of the bubble represents the *p*-value (-log10). (I) SPR sensorgram and steady-state curve for human PTPRU-ECD (analyte) binding to human OSTN (ligand). (J) Summary of SPR results (Figure S3B) testing binding of OSTN (analyte) binding to R2A, R2B and R3 RPTP subfamily members (ligands). (K) Multivariate heatmap for OSTN calculated from normal tissue mRNA expression correlations. (L) Hierarchical 2-way clustering heatmap of normal tissue mRNA expression for OSTN and PTPRU. (M) Schematic representation of the domain architectures of RPTP subfamilies.

**Table 1.** Summary of new Protein Interactions tested in this study

| **Interaction** | **Screen** | **Hit** | **Library** | **Assay** |
| --- | --- | --- | --- | --- |
| GAS1-PTPRA | GAS1 | PTPRA | TM1 | SPR, CSS |
| OMG-PTPRD | OMG | PTPRD | TM1 | SPR |
| OMG-PTPRF | OMG | PTPRF | TM1 | SPR |
| OMG-PTPRS | OMG | PTPRS | TM1 | SPR |
| OMG-PTPRU | OMG | PTPRU | TM1 | SPR |
| OSTN-PTPRU | OSTN | PTPRU | TM1 | SPR |
| MK-RNF167 | MK | RNF167 | TM1 | SPR |
| PTN-RNF167 | PTN | RNF167 | TM1 | SPR |
| PTN-ROR1 | PTN | ROR1 | TM1 | SPR |
| PTN-MFRP | PTN | MFRP | TM1 | SPR |
| SMOC1-SMOC1 | SMOC1 | SMOC1 | TM1 | SPR |
| SMOC1-SMOC2 | SMOC1 | SMOC2 | TM1 | SPR |
| TAFA2-KIR3DL1 | TAFA2 | KIR3DL1 | TM1 | SPR, CSS |
| TAFA2-KIR3DL2 | TAFA2 | KIR3DL2 | TM1 | SPR, CSS |
| TAFA2-KIR3DL3 | TAFA2 | KIR3DL3 | TM1 | CSS |
| TAFA2-KIR2DL5A | TAFA2 | KIR2DL5A | TM1 | CSS |
| TAFA2-RNF167 | TAFA2 | RNF167 | TM1 | SPR |
| TAFA5-RNF167 | TAFA5 | RNF167 | TM1 | SPR |
| LY6H-CD36 | LY6H | CD36 | TM2+ | CSS |
| NRN1-CD36 | NRN1 | CD36 | TM2+ | CSS |
| SCRG1-CD36 | SCRG1 | CD36 | TM2+ | CSS |
| VWC2L-CD36 | VWC2L | CD36 | TM2+ | CSS |

SPR, Surface Plasmon Resonance; CSS, Cell Surface Staining

In a first pass analysis we selected the validated hits of the CRISPRa enrichment screens and performed database searches to calculate overlaps between our screening results and the aggregate of BioGrid, BioPlex, and STRING databases (physical interactions; Membrane and secreted proteome). We observed no overlap between any of these databases and the hits reported in this screen (Figure 2B). As we previously reported for interactome screens of Drosophila and human cell surface proteins (Özkan et al., 2013a; Wojtowicz et al., 2020), high-throughput PPI analysis methods such as Y2H and AP/MS generate mostly false positive interactions for secreted and membrane proteins and are unable to identify genuine interactions found through ELISA and/or cell-based screening methods.

### Oligodendrocyte-myelin glycoprotein (OMG) binds to multiple receptor tyrosine phosphatases

Protein tyrosine phosphorylation is a fundamental regulatory step in intracellular signal transduction and is orchestrated in a coordinated fashion by activities of protein tyrosine kinases (PTKs) and phosphatases (PTPs). PTPs play essential roles in the regulation of growth, differentiation, oncogenic transformation, and other processes (Julien et al., 2010). The classical PTPs include cytoplasmic PTPs and transmembrane receptor protein tyrosine phosphatases (RPTPs), which can be classified into distinct subfamilies according to their domain architecture.

Most RPTPs display features of cell-adhesion molecules (CAMs) with a domain repertoire including MAM (meprin, A-5) domains, Ig (immunoglobulin-like) domains, and FN (Fibronectin) Type III repeats in their extracellular segments (Figure 3M) (Tonks, 2006). In our human in vitro interactome screen, we identified new cell surface binding partners for multiple RPTPs that are likely to mediate cell-cell and/or cell–matrix interactions (Wojtowicz et al., 2020).

In our CRISPRa screen with OMG, we observed the Type R2B subfamily member PTPRU as the top-ranking hit (Figure 3A) with a positive enrichment trajectory over all 3 rounds of selection (Figure 3B). We confirmed binding of OMG to PTPRU by SPR, with a K_D_ of ∼2 µM (Figure 3C). We also identified two members of the R2A subfamily, PTPRF and PTPRS, as well as the R4 subfamily member PTPRA as enriched in the OMG screen (Figure 3D). Type R2A (PTPRD, PTPRF, PTPRS), R2B (PTPRK, PTPRM, PTPRT, PTPRU) and R3 (PTPRB, PTPRH, PTPRJ, PTPRO, PTPRP) are the largest RPTP subfamilies. They all have large ECDs that include FN-III repeats. R2A RPTPs also have Ig domains, and R2B RPTPs have both Ig and MAM domains (Figure 3M). PPIs often occur between phylogenetically related proteins both within and between subfamilies. We examined binding of OMG to all R2A and R2B subfamily members as well as PRPRJ (R3) by SPR. Binding in the micromolar affinity range was observed for all three R2A RPTPs (PTPRD, PTPRF, PTPRS) but only for PTPRU among R2B RPTPs (Figure 3E; S3A). Hierarchical clustering by healthy tissue expression correlations may infer functionally related communities. We therefore examined healthy tissue mRNA expression profiles for OMG, R2A, R2B and R3 RPTP family members from the Human Protein Atlas (Karlsson et al., 2021) and performed a multivariate clustering analysis. OMG clustered with several RPTP family members including binding partners PTPRU and PTPRD (Figure 3F). In the nervous system, these RPTPs are expressed primarily in neurons, and could function as receptors for OMG, which is expressed in oligodendrocytes and some neurons (Figure S3B).

### PTPRU binds to Osteocrin, a primate-specific brain ligand

PTPRU was also identified as a potential hit in a screen for Osteocrin (OSTN). Although our initial ESP ranking showed PTPRJ, a RPTP member of the R3 subfamily, as the top ranking hit for OSTN (Figure 3G), analyzing the enrichment trajectories over all 3 rounds of selections revealed that PTPRJ actually followed a negative trajectory (Figure 3H). By contrast, the R2B family member PTPRU showed a significant positive enrichment trajectory over the course of the screening workflow compared to PTPRJ (Figure 3H). We therefore analyzed binding of OSTN to a panel of R2A, R2B and R3 RPTP members and found that OSTN exclusively bound to PTPRU, with a K_D_ of ∼46 nM (Figure 3I, J, S3C).

OSTN (musclin) is a 130 aa peptide hormone that was originally identified in mouse bone and muscle. It regulates bone growth, supports physical endurance and mediates diverse cardiac benefits of physical activity (Subbotina et al., 2015). These actions could be mediated through OSTN’s binding to the natriuretic peptide clearance receptor (NPR-C) (Moffatt et al., 2007). By binding to NPR-C, OSTN decreases clearance of natriuretic peptides and thereby increases signaling through the NPR-A and NPR-B receptors. In primates, however, the OSTN gene has acquired neuron-specific regulatory elements, and primate OSTN is expressed in cortical neurons and is induced by depolarization in in vitro cultures and by sensory stimuli in vivo. OSTN restricts dendritic growth after depolarization. OSTN expression peaks during the onset of synaptogenesis in fetal development, but it continues to be expressed in neocortex in adults (Ataman et al., 2016). A pairwise correlation of normal human tissue mRNA expression data for OSTN and the R2A, R2B and R3 RPTP subfamilies showed correlation only with PTPRU and its close relative PTPRT (Figure 3K). A hierarchical cluster analysis of human tissue mRNA expression data shows a strong correlation of OSTN and PTPRU in brain and skeletal muscle (Figure 3L; Cluster 3).

### The Growth Arrest Specific 1 (GAS1) protein binds to PTPRA

PTPRA was identified as the highest ranking hit in the GAS1 screen, with the highest ESP score and a consistent positive enrichment over three rounds of positive selection (Figure 4A, B), followed by lower scoring PTPRU and PTPRJ with low enrichment (Figure 4C). PTPRA is a member of the R4 RPTP subfamily (Figure 3M), which have short, highly glycosylated ECDs. GAS1 bound exclusively to PTPRA, with a K_D_ of ∼1 µM (Figure 4D). We also showed that tetramerized GAS1 (GAS1:SA647) exhibits increased binding to K562 cells that overexpress PTPRA, demonstrating that GAS1 is a soluble ligand for cell-surface PTPRA (Figure 4E). No binding was observed to PTPRU or PTPRJ (Figure S4A) and multivariate clustering showed that GAS1 is clustering more closely to PTPRA than to PTPRU or PTPRJ (Figure 4F).

**Figure 4.**
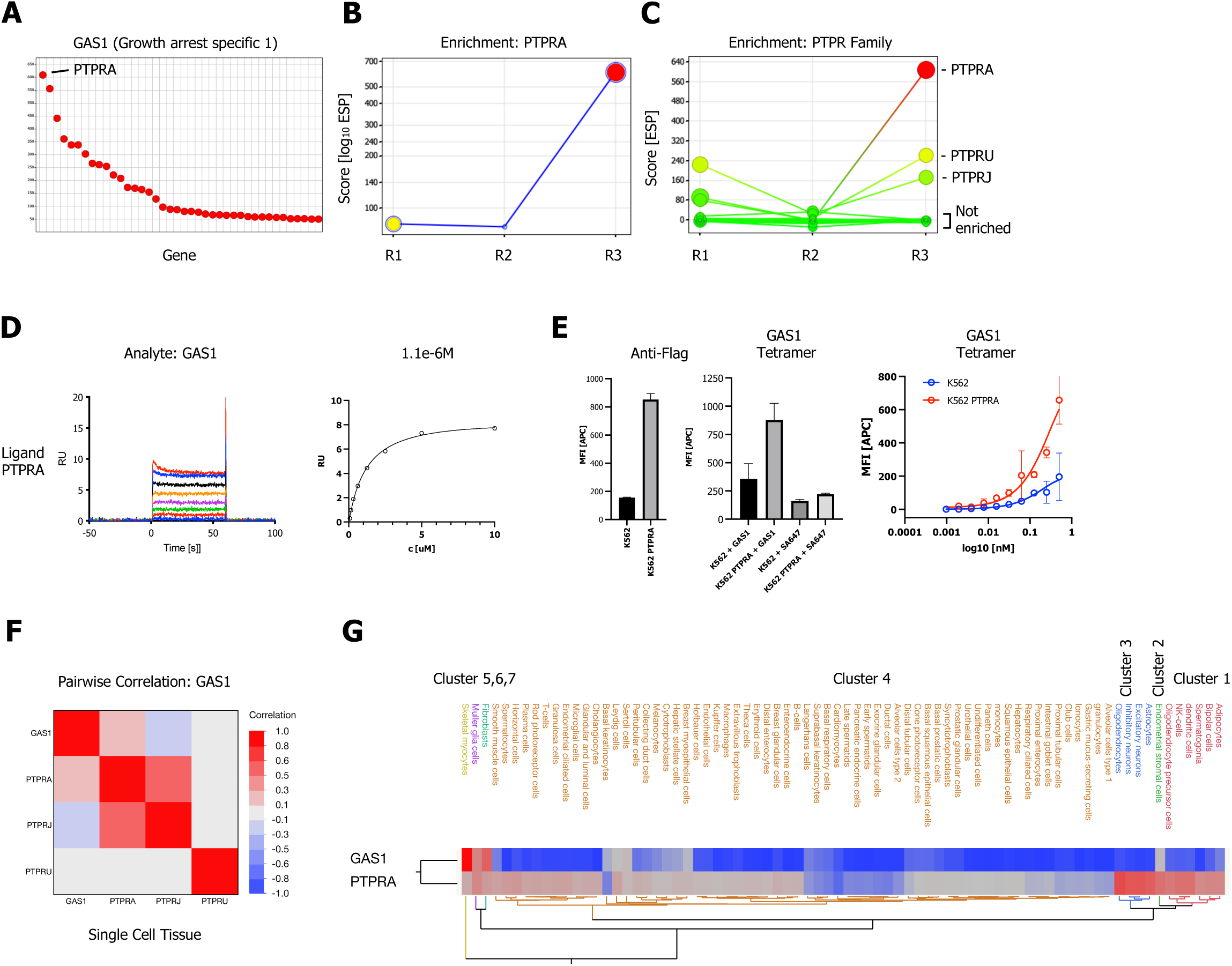
GAS1 CRISPRa enrichment screen identifies RPTP subfamily member PTPRA. (A) Ranked x/y scatter plot for the GAS1 CRISPRa enrichment screen (ESP scores). (B) Trajectory plot of the highest ranking candidate PTPRA for all 3 consecutive rounds of selections in a x/y enrichment plot, size of the bubble represents the p-Value (-log10). (C) Comparison of ESP trajectories for PTPRA and 2 lower scoring RPTP subfamily members (PTPRU, PTPRJ). (D) SPR sensorgram and steady-state curve for human GAS1 (ligand) binding to PTPRA-ECD (analyte) binding in comparison to PTPRU and PTPRJ (Figure S4A; no binding observed). (E) Cell surface staining of K562 cells lenti-virally transduced with FLAG-tagged full-length PTPRA and control K562 cells with GAS1:SA-647 tetramers (400 nM) and analysis by flow cytometry. Data are represented as mean ± SD. (F) Multivariate heatmaps for GAS1 and the PTPRA, PTPRU and PTPRJ calculated from single cell normal tissue mRNA expression correlations. (G) Hierarchical 2-way clustering heatmap of single cell normal tissue mRNA expression for GAS1 and PTPRA shows.

PTPRA is ubiquitously expressed, while GAS1 has a more restricted expression pattern (Figure 4G). GAS1 and PTPRA are both involved in RET tyrosine kinase signaling, as well as in other signaling pathways. GAS1 is related to the GFR1 family of transmembrane proteins, which are coreceptors for the RET receptor tyrosine kinase (RTK). RET-GFR1 complexes bind to glial-derived neurotrophic factor (GDNF), leading to RET autophosphorylation and activation of downstream Akt and MAPK signaling pathways. GAS1 interacts directly with RET and recruits it to lipid rafts. GAS1 binding causes a reduction in GDNF-induced Akt phosphorylation, suggesting that it is a negative regulator of RET signaling. PTPRA also associates with RET signaling complexes and can directly dephosphorylate RET, causing inhibition of RET signaling. PTPRA has not been demonstrated to directly bind to RET, however, and a linkage between PTPRA and RET might be provided by GAS1.

We also examined data from the Cancer Genome Atlas (TCGA; (http://www.cbioportal.org/), and found that in most tumor types high expression of GAS1 is correlated with negative outcomes, while PTPRA expression is associated with favorable outcomes (Figure S4B).

### TAFA-2 selectively interacts with inhibitory Killer-Cell Immunoglobulin-like Receptors (KIRs)

A CRISPRa enrichment screen with TAFA-2 (FAM19A2; chemokine like family member 2) identified two inhibitory killer immunoglobulin-like receptors (KIRs), KIR3DL1 and KIR3DL3, which are selectively expressed on natural killer (NK) cells, as the highest ranking hits by ESP scoring (Figure 5A). KIRs are a polymorphic subfamily of MHC class I receptors (Li and Mariuzza, 2014; Pende et al., 2019; Sivori et al., 2019). KIR3s have D0, D1, and D2 Ig-like domains. KIR2s have only D1 and D2 domains, except for KIR2DL5, which has a D0 and D2 but lacks D1 (Figure 5C). The structure of KIR3DL1 complexed to an HLA-B reveals that the helices and bound peptide of the HLA engage with the D1 and D2 domains of the KIR, while the D0 domain extends down toward the b-2-microglobulin subunit and engages sequences that are highly conserved among all HLA A and B alleles (Li and Mariuzza, 2014).

**Figure 5.**
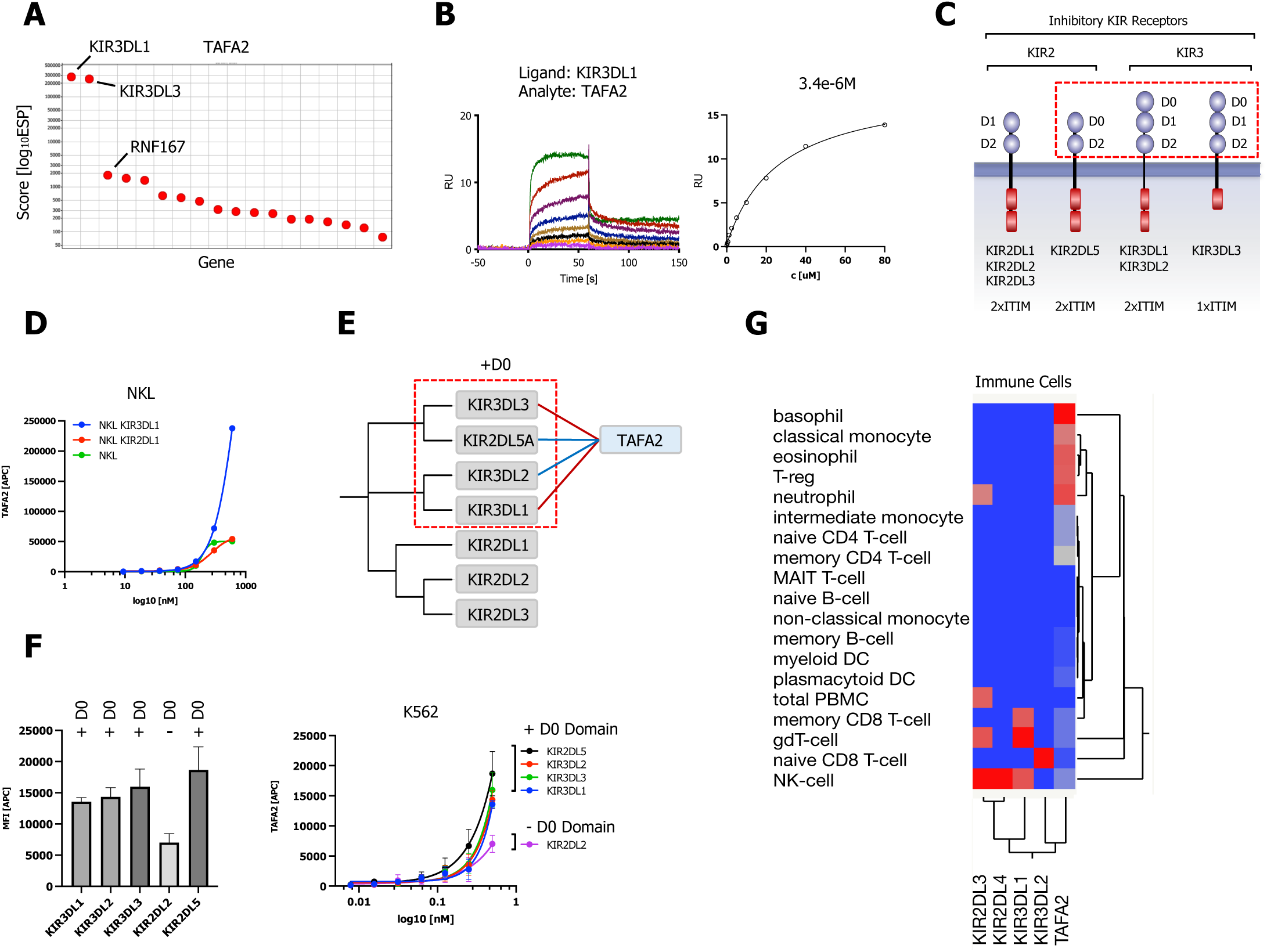
KIR subfamily PPIs Identified by CRISPRa screening and PHA approach for TAFA2. (A) ESP rank plot of the TAFA2 CRISPRa enrichment screen. (B) SPR sensorgrams and steady-state curves for human TAFA2 (analyte) binding to KIR3DL1-ECD. (C) Schematic representation of the domain architecture of KIR2 and KIR3 subfamily of inhibitory KIR receptors. (D) Cell surface staining of NKL cells expressing KIR3DL1, KIR2DL1 or NKL control cells with TAFA2:SA-647 tetramers (200 nM) and analysis by flow cytometry. (E) Dendrogram of the KIR2 and KIR3 subfamily calculated from multiple sequence alignments of KIR receptor ECDs and PPIs observed in the CRISPRa screen (red) and predicted by PHA (blue). (F) Cell surface staining of K562 control cells or K562 cells lenti-virally transduced with full-length KIR3DL1, KIR3DL2, KIR3DL3, KIR2DL2 or KIR2DL5A (FLAG-tagged; Figure S5C) with TAFA2:SA-647 tetramers (200 nM) and analysis by flow cytometry. Data are represented as mean ± SD. (G) Hierarchical 2-way clustering heatmap of immune cell mRNA expression data for TAFA2, KIR2 and KIR3 subfamily members.

**Figure 6.**
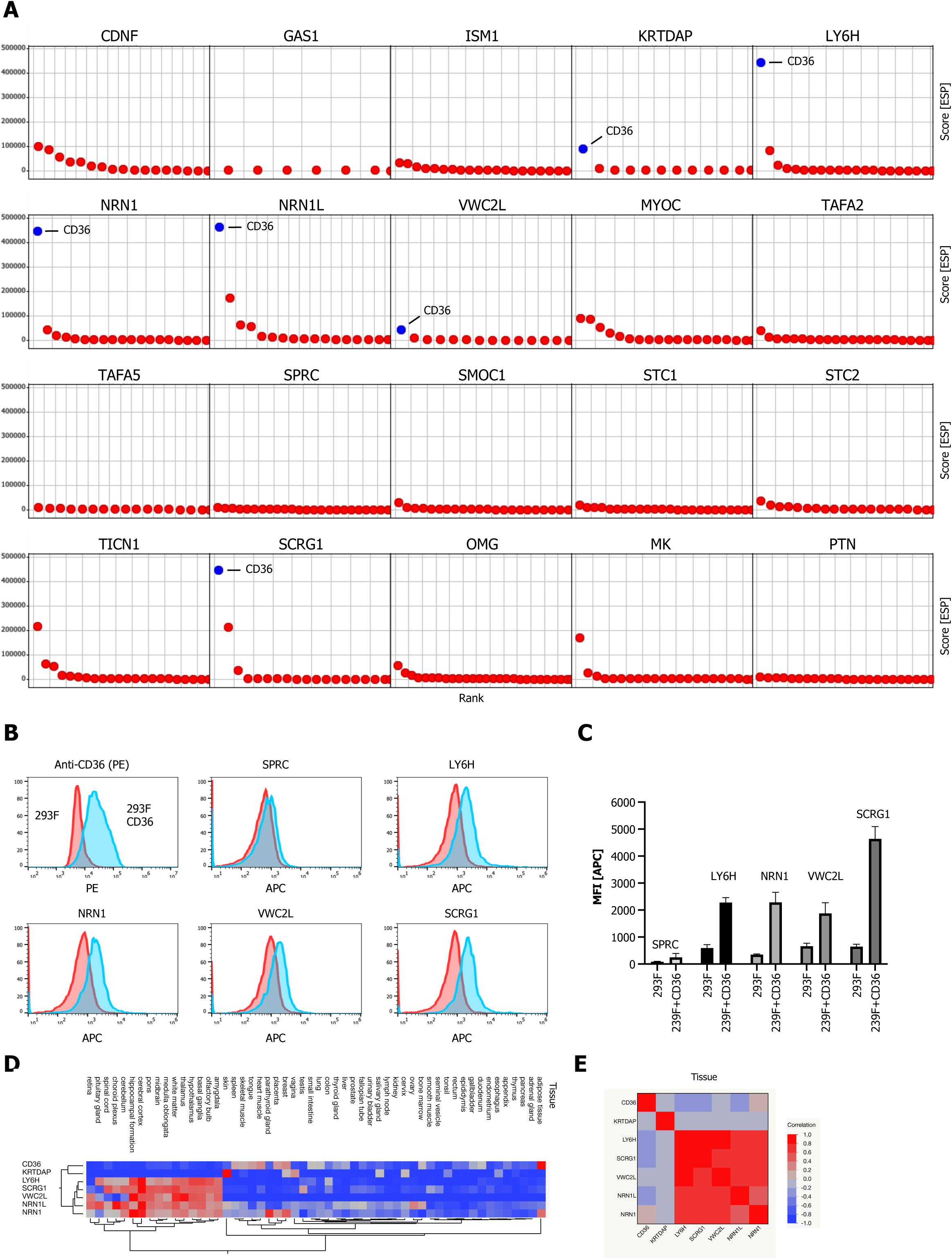
Multifunctional scavenger receptor CD36 binds multiple secreted ligands. (A) Ranked ESP scatter trellis plots of all analyzed TM2+ library screens shows CD36 as the highest ranking hit for multiple screens (CD36 indicated in blue). (B-C) Cell surface staining of full-length CD36-transfected and control cells with SA-647 tetramerized (400 nM) LY6H, NRN1, VWC2L or SCRG1 and analysis by flow cytometry, SPRC tetramers (SPRC:SA647) served as a negative (non CD36 enriched) control. (B) representative histograms (red, control cells; blue, CD36 positive cells). (C) Quantification of flow data (n=3); data are represented as mean ± SD. (D-E) Hierarchical 2-way clustering heatmap and multivariate correlation of normal tissue cell mRNA expression data for CD36 and the indicated CD36 enriched screens.

We observed binding of TAFA-2 to KIR3DL1 to by SPR, with a K_D_ of ∼33 µM (Figure 5B, S5A). Binding to KIR3DL3 was at the limit of detection and did not saturate (Figure S5B). To examine binding at the cell surface, fluorescent TAFA-2 tetramers (TAFA2:SA647) were incubated with NKL cells expressing either full-length KIR3DL1 or KIR2DL1. Flow cytometry analysis revealed concentration-dependent binding of TAFA-2 to cells expressing KIR3DL1, but not to those expressing KIR2DL1 (Figure 5D). To further define KIR binding specificity, we tested binding of TAFA-2 tetramers (TAFA2:SA647) to K562 cells expressing KIR3DL1, KIR3DL2, KIR3DL3, KIR2DL2 or KIR2DL5A. We observed higher concentration-dependent binding of TAFA-2 to cells expressing the KIR3s or KIR2DL5A, which all have D0 domains, compared to to KIR2DL2, which does not have a D0 domain (Figure 5E, F). These data suggest that D0 domains are required for TAFA-2 binding.

TAFA-2 is a member of a highly conserved 5-gene family (TAFA1-5) of chemokine-like peptides (neurokines) expressed in the brain. Like other chemokines, TAFAs 1, 4, and 5 bind to G protein-coupled receptors (GPCRs). TAFAs 1-4 all complex with neurexins during their passage through the ER/Golgi pathway, leading to formation of disulfide-bonded cell-surface neurexin-TAFA complexes (Khalaj et al., 2020; Sarver et al., 2021; Tom Tang et al., 2004). Although TAFA-2 has only been examined in the brain, the gene is also expressed in the immune system. Its expression is restricted to naïve and memory regulatory T cells (T-regs), basophils, and neutrophils, with the highest expression levels being observed in basophils (Figure 5G). Neurexins are not expressed in these cell types, so TAFA-2 may be secreted as a monomer or complexed to another protein. The observed interactions of TAFA-2 with D0 domains of KIRs suggest that expression of the chemokine by regulatory T cells or basophils might modulate KIR signaling in NK cells in response to binding of HLA on target cells.

### The Scavenger Receptor CD36 acts as a receptor for a broad range of secreted ligands

CD36, also known as SCARB3 or glycoprotein 4 (GPIV), is a multifunctional Type B scavenger receptor with two transmembrane domains and a ∼410 aa spanning extracellular domain. CD36 is known to bind to many ligands (Silverstein and Febbraio, 2009). In our analysis of TM2+ library screens we identified CD36 as a top hit in several screens: KRTDAP, LY6H, NRN1, NRN1L, VWC2L and SCRG1 (Figure 6A). To examine potential binding of these secreted ligands to CD36 on the cell surface, fluorescent tetramers of LY6H, NRN1, VW2CL and SCRG1 were incubated with 293F cells or 293F cells expressing full-length CD36 (Figure 6B). We observed binding of LY6H, NRN1, VW2CL and SCRG1 to CD36 by FACS. SPRC, which showed no enrichment of CD36 in its screen, served as a control and did not bind to CD36 (Figure 6B, C). All CD36 enriched secreted ligands (LY6H, NRN1, NRN1L, VWC2L, SCRG1) except for KRTDAP show a strong correlation and cluster in brain tissue, which expresses CD36 mRNA only at low levels (Figure 6D). However, CD36 protein is expressed in brain microglia, and CD36-mediated debris uptake regulates brain inflammation in neurodegenerative disease models (Dobri et al., 2021; Grajchen et al., 2020).

### Other Screens

In Figure S6-S7, we show results of additional screens: Midkine (MK), Pleiotrophin (PTN), TAFA-2, and TAFA-5. We analyzed the top 20 highest ranking candidates for MK, PTN, TAFA-2 and TAFA5 and found a strong overlap for several candidates (Figure S6A). We identified the transmembrane PA-TM-RING E3 ligase RNF167 as a high ranking hit shared between screens for PTN, TAFA-2 and TAFA-5. PTN and MK are two members of the neurite growth-promoting factor family. Like TAFA-2, TAFA-5 is a member of the TAFA superfamily and is highly expressed in in the brain tissue, especially hippocampus, cerebral cortex, white matter and ganglia (Figure S6F). *TAFA5* knockout mice display increased depressive-like behaviors and impaired hippocampus-dependent spatial memory (Huang et al., 2021). We observed binding of RNF167 to PTN and MK as well as TAFA-2 and TAFA-5 by SPR (Figure S6B, C). RNF167 is a member of the PA-TM-RING E3 ligase family, which has approximately 20 members. These E3 ligases are minimally defined by three conserved domains, a protease-associated (PA) domain that acts as a substrate recruitment domain, a transmembrane domain (TM), and a RING-H2 finger (RNF) (Nakamura, 2011). RNF167 (also known as Godzilla) is widely expressed in all tissues including brain tissue and has recently been implicated in the regulation of the AMPA receptor (AMPAR) (Ghilarducci et al., 2021; Lussier et al., 2012). 2-way hierarchical clustering shows a high correlation between RNF167, TAFA-2, TAFA-2 and PTN in brain tissue (Figure S6F).

The transmembrane RTK receptor ROR1 and the Membrane Frizzled-related protein MFRP were identified as shared hits among several screens (PTN, MK, TAFA-2, TAFA-5). We observed binding of PTN to ROR1 and MFRP (Figure S6D). Intestingly, both receptors share a evolutionary conserved Frizzled domain in the ECD (Yan et al., 2014). ROR1 and its closely related paralog ROR2 are receptors for Wnt5a and other Wnts in the planar cell polarity (PCP) pathway (Endo et al., 2022; Green et al., 2014; Minami et al., 2010). MFRP (Membrane frizzled-related protein) is type II transmembrane protein with an extracellular FZD domain. MFRP is predominantly expressed in the retinal pigment epithelium (RPE) with high expression in brain tissue (Choroid plexus and required for both prenatal ocular growth and postnatal emmetropization (Katoh, 2001; Sundin et al., 2008).

SMOC1, a SPARC-related ligand overexpressed in brain tumors (Brellier et al., 2011) recovered BST2, IGSF23 and SMOC2 as the highest ranking hits (Figure S7A,B). We only observed binding of SMOC1 to SMOC and SMOC1 by SPR (Figure S7C). SMOC2 is a matricellular protein which promotes matrix assembly and is involved in endothelial cell proliferation and migration, more recently SMOC2 () variants were also reported to be play a role in BMP signaling (Long et al., 2021).

## DISCUSSION

In an effort to accelerate the discovery of novel interactions between secreted ligands and the membrane proteome, we have developed a high-throughput cell based CRISPRa enrichment screening workflow by employing customized, pooled cell surface receptor sgRNA libraries in combination with MACS to enrich for receptor-expressing cells. We defined a list of 80 high-priority secreted ligands that are likely to have receptors that have not been previously identified. 20 of these were successfully expressed, biotinylated, coupled to streptavidin magnetic beads, and used for enrichment of CRISPRa mediated receptor-expressing cells via MACS. We identified the receptors that are enriched in the isolated cells via deep sequencing of their sgRNAs (Figures 1, 2).

To validate the results of the screens, we used SPR and cell binding methods to prove that the receptor candidates actually bound to the ligands used for screening. We then expanded the set of receptor candidates by taking advantage of homology, examining other members of the gene families identified in the initial screens for binding. Using these approaches, we identified 14 candidate receptors for the 12 screened ligands. In some cases (OMG, TAFA-2), a single ligand bound to multiple receptors in the same family, while in other cases (CD36, RNF167), a single receptor bound to several unrelated ligands (Figure 2C). Our results highlight the inability of standard high-throughput in vitro screening methods to identify genuine ECD interactions, because none of the hits from our screen were found in protein interaction databases (Figure 2B). This was previously observed when the results of ELISA-based in vitro screens using multimeric ECD fusion proteins were compared with PPIs represented in the databases (Martinez-Martin, 2017; Özkan et al., 2013a; Ranaivoson et al., 2019; Söllner and Wright, 2009; Taouji et al., 2009; Verschueren et al., 2020; Wojtowicz et al., 2020).

Our results suggest that the cell-based screening strategy is primarily limited by two factors. First, fewer than 1/3 of the high priority ligand candidates were expressed at high enough levels to allow purification for screening. If it were possible to express all of the 206 ligands we identified as targets, one might expect to be able to identify receptors for more than 100 of them. However, it is unclear how this might be accomplished. Second, the TM2+ screen yielded a lower rate of enriched candidates than the TM1 screen, identifying only the scavenger receptor CD36 (Figure 2C). Many TM2+ receptors are protein complexes, and in these cases sgRNA-driven expression of single genes would not be able to generate cells that make functional receptors, unless the parental cell K562 line also made the other receptor complex components. Also, expression of TM2+ receptors may often require chaperones that might not be present in K562 cells.

### New biology revealed by the screen

The most striking finding from our screen was the identification of three new ligands for RPTPs, a diverse set of cell signaling receptors whose functions are less well understood than those of RTKs. GAS1, a protein involved in control of cell growth, binds to PTPRA, an R4 RPTP that has no known ligand (Figure 4). PTPRA is a ubiquitously expressed signaling protein that both positively and negatively regulates tyrosine kinase signaling via dephosphorylation of Src and of RTKs. GAS1 may link PTPRA to RET RTK complexes. OMG, a ligand expressed by oligodendrocytes and some neurons, binds to all three R3 RPTPs and to one of the R2B RPTPs, PTPRU (Figure 3). All of these RPTPs are expressed in the brain. Most interestingly, OSTN, a hormone that regulates bone and muscle growth in rodents but has acquired brain-specific expression and function in primates (Ataman et al., 2016) binds only to PTPRU (Figure 3). OSTN was known to be a ligand for NPR-C, which regulates the levels of natriuretic peptides, but this is unlikely to explain its function in the primate brain. PTPRU, however, is expressed in the brain and is closely related to the three other R2B RPTPs, which are homophilic adhesion molecules that regulate cadherin-mediated cell adhesion. Interestingly, PTPRU cannot mediate cell adhesion on its own, and is thought to lack phosphatase activity (Hay et al., 2020), so it may have a distinct function from the other R2B RPTPs.

We also found an unexpected association between TAFA-2, which is known as a neurokine (brain chemokine), and KIRs, which are inhibitory MHC class I receptors that are expressed only by NK cells (Figure 5). The TAFA2 gene is also expressed by regulatory T cells and basophils, and TAFA-2 might be a soluble ligand that modulates NK signaling in response to MHC class I engagement.

In summary, we implemented a proteo-genomic screening workflow, combining CRISPRa pooled cell surface libraries with a MACS enrichment strategy, in order to accelerate the identification of interactions between the secreted and membrane proteomes. We report new receptor-secreted ligand PPIs that are potentially involved in a wide variety of signaling processes. Implementation of cell-based screening strategies based on our approaches might allow elucidation of receptor-ligand relationships for many proteins that are currently orphans, and this has the potential to identify therapeutic targets and define new biological processes.

## MATERIAL AND METHODS

### Curation of the human membrane and secreted proteome and selection of secreted bait proteins

Two lists of human membrane and secreted proteins were generated using the Human Protein Atlas (HPA) database (https://www.proteinatlas.org/humanproteome/tissue/secretome). Lists were checked for overlap and master lists were generated using the HPA majority decision-based method (MDM). Metadata for all master list proteins was extracted from UniProt https://www.uniprot.org and sequence information was validated by the SIgnalP-5.0 (http://www.cbs.dtu.dk/services/SignalP/) (Almagro Armenteros et al., 2019), TMHMM-2.0 (https://services.healthtech.dtu.dk/service.php?TMHMM-2.0) and PredGPI (http://gpcr.biocomp.unibo.it/predgpi/) (Pierleoni et al., 2008) prediction servers. Following curation, canonical protein sequences were extracted from UniProt and compiled for back translation and optimization by GeneArt/Life Sciences Technology for gene synthesis.

### Database integration and hierarchical clustering analysis

Interaction datasets were downloaded from BioGRID (https://thebiogrid.org, 4.4.210, physical interactions), Bioplex (https://bioplex.hms.harvard.edu, Hek293 v3.0 and HCT116 v1.0) and STRING (https://string-db.org/; v11.5; physical dataset; interaction score > 0.4). To calculate the overlap between all obtained datasets and out own study, interactions were restricted to physical interactions reported and a Venn diagram was visualized (Heberle et al., 2015). Tissue expression datasets (normal tissue and TCGA) were downloaded from the The Human Protein Atlas (https://www.proteinatlas.org; v21.1). Unsupervised hierarchical clustering of normalized mRNA gene expression by tissue was performed with Ward linkage and correlation distance were plotted as heatmaps using JMP Pro (v16).

### Generation of secreted mammalian expression plasmids

Genes encoding curated secreted proteins of interest (SPOI) were synthesized at GeneArt/Life Sciences Technologies and subcloned into pD649-SPOI-AviTag-6xHis. Genes were subcloned in-frame with the endogenous signal peptide and downstream AviTag-6xHis modules via 5’ NheI and 3’ AscI sites. A MaxiPrep of plasmid DNA was provided at 1 ug/ml in 20 mM Tris, pH 8.0. For complete plasmid sequences of all 80 SPOI bait expression vectors, see Data Sx. All Plasmids (80) will be made available through Addgene.

### Generation of Expression Plasmids for Full-Length Proteins

Genes encoding full-length proteins were synthesized at GeneArt/Life Sciences Technologies and subcloned into the pHR expression vector. Plasmids contain a kozak sequence, HA signal peptide, a FLAG tag (to facilitate cell-surface expression analysis) and the remaining full-length coding region of the gene, followed by a stop codon. Synthesized DNA were subcloned into pHR-kozak-SPHA-1xFlag using using AscI and BamHI sites. A MaxiPrep of plasmid DNA was provided at 1 ug/ml in 20 mM Tris, pH 8.0. For transgene sequences, see Data Sx. Plasmids will be made available through Addgene.

### Generation of pDisplay Plasmids

Genes encoding curated secreted proteins of interest (SPOI) were subcloned in frame into pHR expression vector with the endogenous signal peptide and a downstream 1xFLAG-CD8-Stalk-CD8-TM-CD8-ICD or 1xHA-CD8-Stalk-CD8-TM-CD8-ICD module via 5’ NheI and 3’ AscI sites.

### Production of purified proteins

Proteins were produced in Expi293F cells using transfection conditions following the manufacturer’s protocol. After harvesting of cell conditioned media, 1 M Tris, pH 8.0 was added to a final concentration of 20 mM. Ni-NTA Agarose (Qiagen) was added to ∼5% conditioned media volume. 1 M sterile PBS, pH 7.2 (Gibco) was added to ∼3X conditioned media volume. The mixture was stirred overnight at 4° C. Ni-NTA agarose beads were collected in a Buchner funnel and washed with ∼300 ml protein wash buffer (30 mM HEPES, pH 7.2, 150 mM NaCl, 20 mM imidazole). Beads were transferred to an Econo-Pak Chromatography column (Bio-Rad) and protein was eluted in 15 ml of elution buffer (30 mM HEPES, pH 7.2, 150 mM NaCl, 200 mM imidazole). Proteins were concentrated using Amicon Ultracel filters (Millipore) and absorbance at 280 nm was measured using a Nanodrop 2000 spectrophotometer (Thermo Fisher Scientific) to determine protein concentration.

### Biotinylation and FPLC Purification

Where indicated, proteins were biotinylated as described previously (Özkan et al., 2013b). Briefly, up to 10 mg of protein was incubated at 4° overnight in 2X Biomix A (0.5 M bicine buffer), 2X Biomix B (100 mM ATP, 100 mM MgOAc, 500 μM d-biotin), Bio200 (500 uM d-biotin) to a final concentration of 20 uM, and 60-80 units BirA ligase in a final volume of 1 ml. Proteins were further purified by size-exclusion chromatography using an S200 Increase or a Superose S6 column (GE Healthcare), depending on protein size, on an ÄKTA Pure FPLC (GE Healthcare), FPLC traces for purified proteins used for the CRISPRa enrichment screens and SPR validation can befound in Figure S8 and S9.

### Lentivirus production

HEK293T (LentiX) cells (female-derived kidney cell line) were grown in DMEM complete media (Thermo Fisher) supplemented with 10% FBS, 2 mM L-glutamine, 50 U ml^−1^ of penicillin and streptomycin, and used to package lentivirus using Fugene HD (Promega) in OptiMem (Thermo) as per the manufacturer’s instructions. Third generation packaging plasmids were used for the pooled sgRNA CSR libraries. After 72h, lentivirus containing media was harvested, filtered (0.45 uM pore, PVDF) and concentrated using PEG-it (SBI) according to the manufacturer’s protocol or lentivirus containing media was used directly to infect the specified cell line.

### CRISPRa enrichment screen

K562 cells stably expressing the sunCAS9 system carrying the pooled sgRNA CSR libraries (TM1; TM2+) were expanded and 50 million cells per screen and library were harvested, washed three times with cold MACS (Miltenyi) buffer and resuspended in 2 ml MACS buffer in a sterile 5 ml Eppendorf tube. Cells were then labeled with magnetic streptavidin microbeads complexed with biotinylated bait protein (50 ul streptavidin microbeads; 1 uM biotinylated protein), mixed and incubated at 4°C for 30 min (tumbling). After labeling, cells were washed twice with cold MACS buffer (300xg, 10 min), resuspended in 1 ml MACS buffer and passed through a 40 uM cell strainer, to obtain a single-cell suspension, directly onto the LS-Column (Miltenyi) for magnetic bead separation according to the manufacturers protocol. Briefly, after applying the labelled cells onto the LS-Column, unlabeled cells pass through and are discarded while labeled cells are retained in the magnetic field, the LS-Column is washed three times with 3 ml of ice cold MACS buffer. For elution of positively selected cells, the column is removed from the separator (magnet) and the magnetically labelled cells are flushed into a 15 ml Falcon tube with fresh media (RPMI complete), washed once, resuspended and transferred to a T25 culture flask for expansion.

### Genomic DNA extraction, library amplification and deep sequencing

Genomic DNA was isolated using QiaAmp DNA Blood Maxi or QiaAmp DNA mini kits (Qiagen) according to the manufacturer’s instructions, genomic DNA was then amplified using Herculase II polymerase (Agilent) as described previously (Deans et al., 2016). To prepare the sgRNA sequencing library, the integrated sgRNA-encoding constructs were PCR amplified using Agilent Herculase II Fusion DNA Polymerase, followed by a second PCR amplification introducing sample-specific Illumina index barcodes and adapters for deep sequencing.

### Cell-Surface binding assay with streptavidin tetramerized secreted ligand

To examine PPIs at the cell surface, we performed cell-surface protein binding assays using K562 or HEK293F cells. K562 cells were used for pre-evaluation of potential base line binding for all secreted proteins used in the CRISPRa screening workflow. HEK293F cells were transfected using Fugene6 according to manufacturer’s protocol (Promega) with expression plasmids encoding full-length proteins containing an N-terminal tag (FLAG or HA) or a pDisplay plasmid encoding the extracellular domain (ECD) of the protein of interest in fusion with a FLAG-Tag or HA-Tag, CD8 hinge and CD8-transmembrane domain. Two days following transfection, cultures were harvested, cells were spun down for 4 min at 1600 rpm (∼400x g), washed twice with cold MACS buffer (Miltenyi) and resuspended to a final density of ∼3 x 10^6^ cells/ml. To generate tetramerized secreted ligands to test for binding to cells expressing full-length proteins, FPLC-purified biotinylated proteins (see above) were incubated with streptavidin tetramers conjugated to Alexa647 Fluor (SA-647) (Thermo Fisher Scientific) at a 4:1 molar ratio on ice for at least 15 minutes. To assess cell-surface expression of full-length or ECD displayed proteins, 1:200 mouse anti-FLAG-647 (CST) or anti-HA antibody (CST) staining of cells was also performed in parallel where indicated. Approximately 150,000 cells were incubated with Protein:SA-647 complexes or antibody in a final volume of 100 ul in 96-well round-bottom plates (Corning) for 1 hour at 4° C protected from light. Following incubation, cells were washed two times with 200 ul cold MACS buffer and resuspended in 200 ul cold MACS buffer with 1:3000 propidium iodide (Thermo Fisher Scientific). Immunofluorescence staining was analyzed using a Cytoflex (Beckman Coulter), and data were collected for 20,000 cells. Data were analyzed using FlowJo v10.4.2 software. All data report mean fluorescence intensity (MFI). Concentration-dependent binding of Protein:SA-647 to full-length receptor-expressing, but not mock control cells, was deemed indicative of cell-surface binding.

### SPR Experiments

SPR experiments were performed using a Biacore T100 instrument (GE Healthcare). FPLC-purified biotinylated proteins (ligands) in HBS-P+ Buffer (GE Healthcare) were captured on a Streptavidin (SA) Series S Sensor Chip (GE Healthcare). Chip capture was performed in HBS-P+ buffer (GE Healthcare) to aim for ∼100-200 ligand response units (RU). Flow cell 1 was left empty to use this flow cell as a reference flow-cell for on-line subtraction of bulk solution refractive index and for evaluation of non-specific binding of analyte to the chip surface using Biacore T100 Control Software (version 3.2) (GE Healthcare). FPLC-purified non-biotinylated protein was used as analyte. Analytes were run in HBS-P+ buffer using 2-fold increasing protein concentrations to generate a series of sensorgrams. A table of all SPR conditions for each ligand-analyte pair tested including concentration range of 2-fold analyte dilutions, injection rate, injection and dissociation times and regeneration conditions can be found in Data S4.

## Acknowledgments

The authors thank the Howard Hughes Medical Institute (K.C.G.), and the G. Harold and Leila Y. Mathers Charitable Foundation (K.C.G.).

## Author Contributions

Conceptualization, K.C.G., D.H.S.; Methodology, D.H.S., M.C.B. and K.G.G.; Library Design: D.H.S., K.Z., G.T.S., M.C.B. and K.C.G.; Screening, D.H.S., L.T.H.; Validation, D.H.S., S.C.W.; Analysis, D.H.S.; Investigation, D.H.S., L.T.H., and K.C.G.; Data Curation, D.H.S., K.Z.; Writing – Original Draft, D.H.S. and K.Z.; Writing – Review & Editing, K.C.G., D.H.S, and K.Z.; Visualization, D.H.S.; Supervision, K.C.G.; Funding Acquisition, K.C.G.

## SUPPLEMENTAL DATA

**Data S1. Secreted protein library**

Information for all 80 proteins including ID, UniProt, Entry name, Protein Name and full-length protein sequence.

**Data S2. Plasmid sequences**

Complete DNA sequences for all 80 secreted bait proteins and full length pD649 expression plasmid sequences.

**Data S3. Pooled CRISPRa library source list**

Information for both customized cell surface libraries, library id, target id and gRNA sequences for the TM1 and TM2+ libraries.

**Data S4. SPR Conditions**

Table of SPR conditions for all ligand-analyte pairs tested including ligand RU, maximum analyte concentration, analyte RU at maximum concentration, injection time (seconds), injection rate (ul/minute), dissociation time (seconds), and regeneration conditions.

---

**Figure S1.**
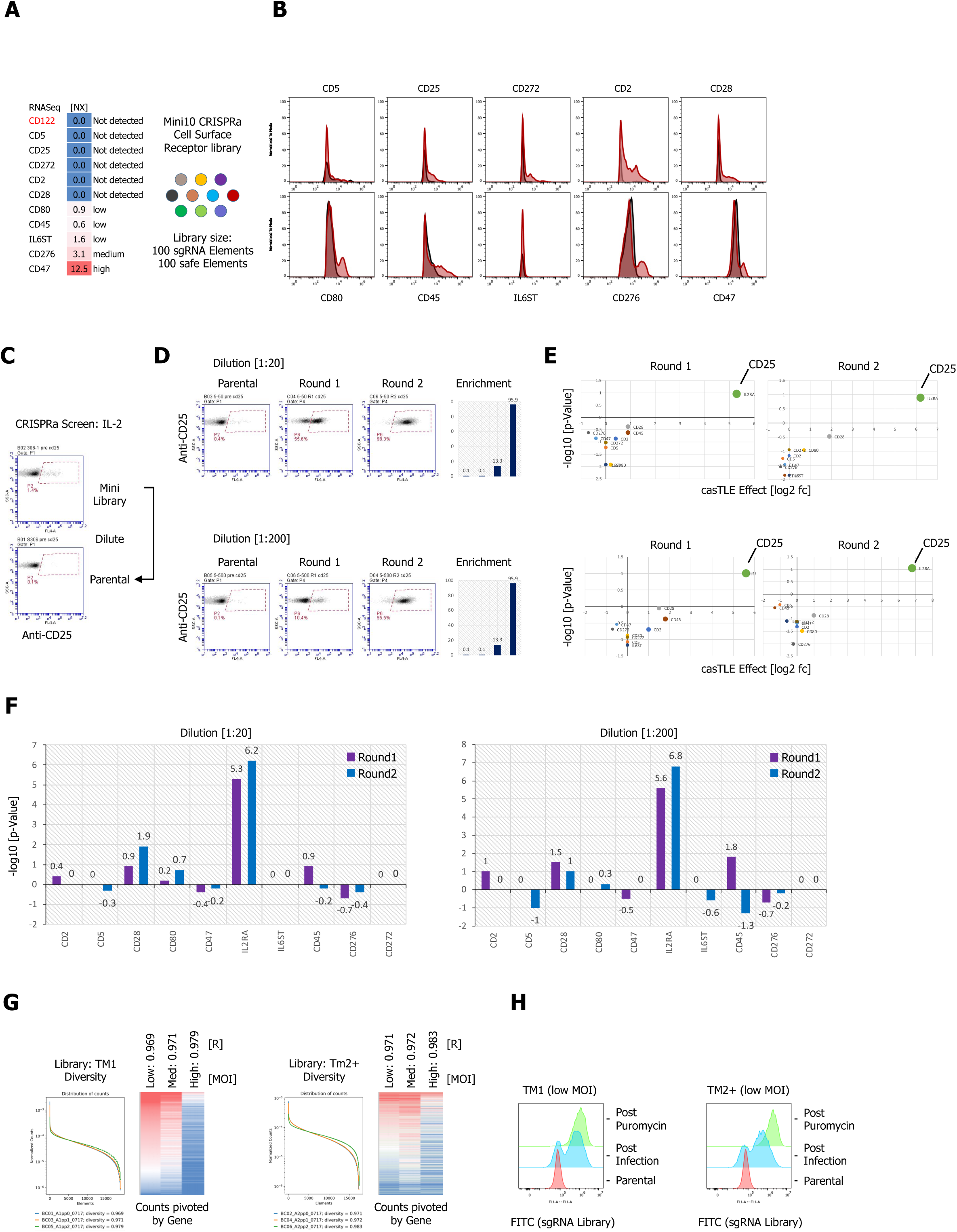
(A) Normalized mRNA expression data of target cell surface receptors selected for a CRISPRa mini-library of 10 genes in K562 cells. (B) Antibody staining of the K562 mini-library after lentiviral infection and puromycin selection in comparison to parental, untransduced K662 cells using APC-labelled primary antibodies. (C-F) Proof of concept CRISPRa enrichment screen using human IL-2 by simulation of a full library screen by diluting the mini-library by factor 20 and 200 and preforming 2 consecutive rounds of magnetic bead selection followed by gDNA extraction, barcoding and deep sequencing. (C) Cell surface staining of the parental and undiluted mini-library using tetramerized IL-2 (IL-2:SA637; 200nM). D) Enrichment over 2 rounds of consecutive magnetic bead selection was detected by tetramer (200nM) staining with human IL-2 (Parental, Round 1 and Round 2) for both simulated libraries post MACS. (E) Visualization of the deep sequencing analysis. Results are visualized by x/y scatter plot: casTLE-Score (log2) / pValue (-log10), size of the hit represents the casTLE-Effect + casTLE-Score. (F) Candidate hits of both rounds of enrichment visualized by bar graphs: casTLE-Score (log2) / pValue (-log10) for each gene in the mini-library. (G) Customized cell surface library quality control. Pooled, customized and target-specific TM1 (∼1800) and TM2+ (∼1750) sgRNA libraries (10 sgRNA per gene together with a safe sgRNA reference pool) were designed, cloned and lentivirally infected into K562-CRISPRa cells at low MOI, medium and high MOI. (H) After puromycin selection genomic DNA was extracted and subjected to deep sequencing for library validation by plotting the library diversity and counts for the 3 MOIs. Right panel: For the final libraries, low MOI libraries for TM1 and TM2+ were analyzed pre and post Puromycin selection by FACS.

**Figure S2.**
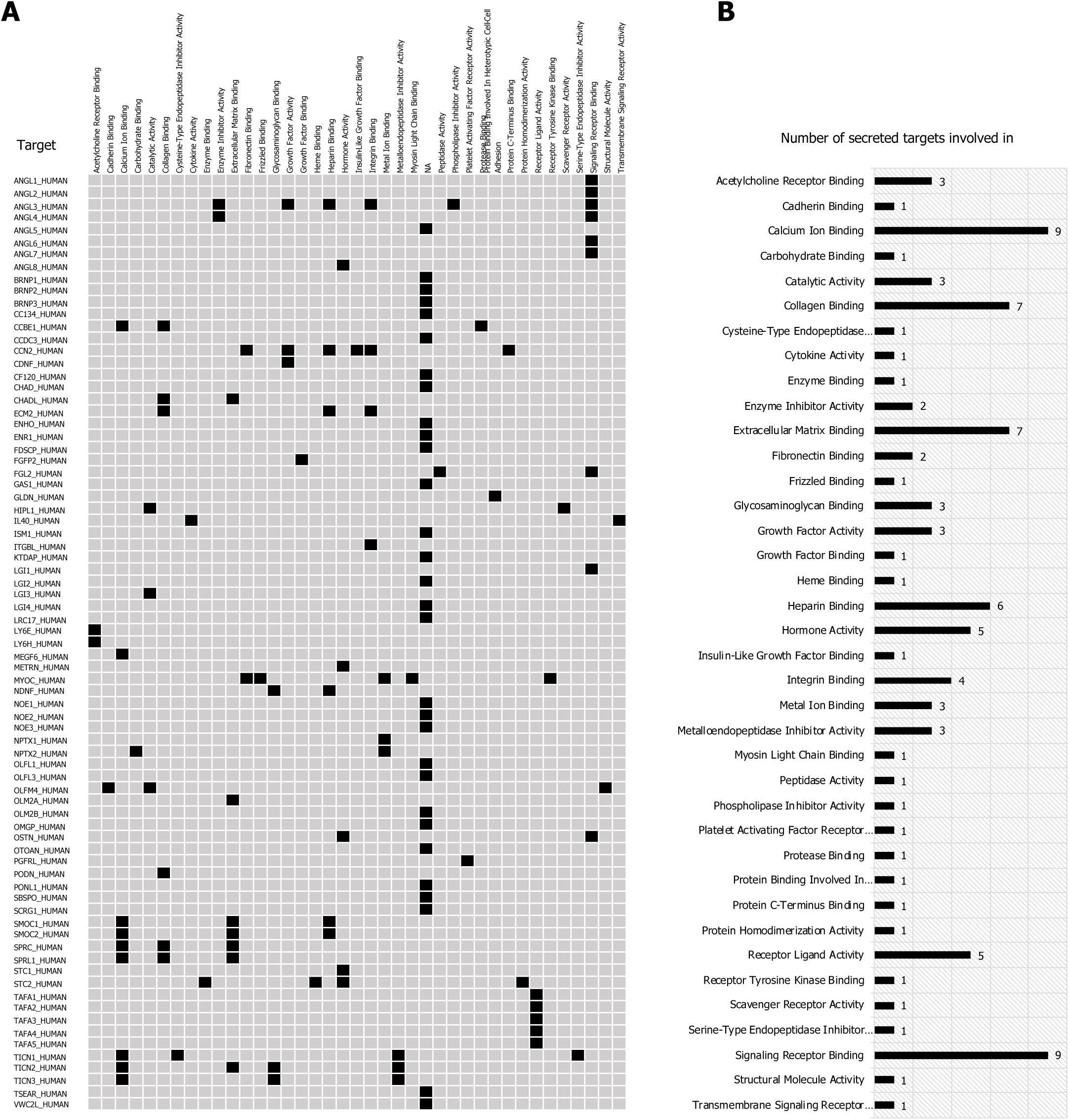
(A) High value secreted orphans (80) classified by molecular function and signaling (B) using GO-term annotations.

**Figure S3.**
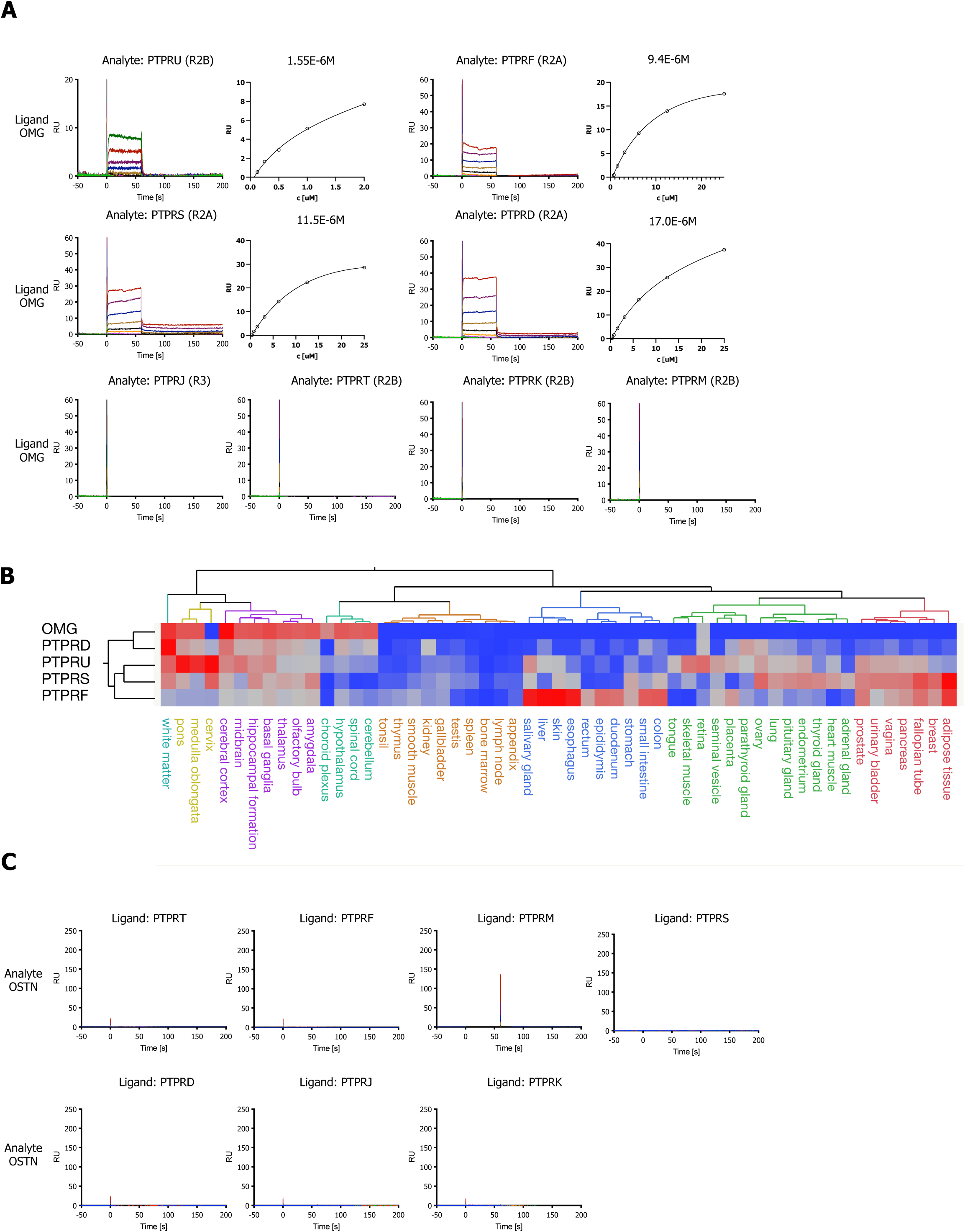
(A) SPR sensorgrams and steady-state curves for human OMG (ligand) and RPTP subfamily members (analytes): PTPRU, PRPRF, PTPRS, PTPRD, PTPRJ, PTPRT, PTPRK, PTPRM (ECDs). (B) Hierarchical 2-way clustering of mRNA expression data from normal tissue for OMG and RPTP subfamily members. (C) SPR sensorgrams curves for human OSTN (analyte) and RPTP subfamily members (ligands): PRPRF, PTPRS, PTPRD, PTPRJ, PTPRT, PTPRK, PTPRM (ECDs).

**Figure S4.**
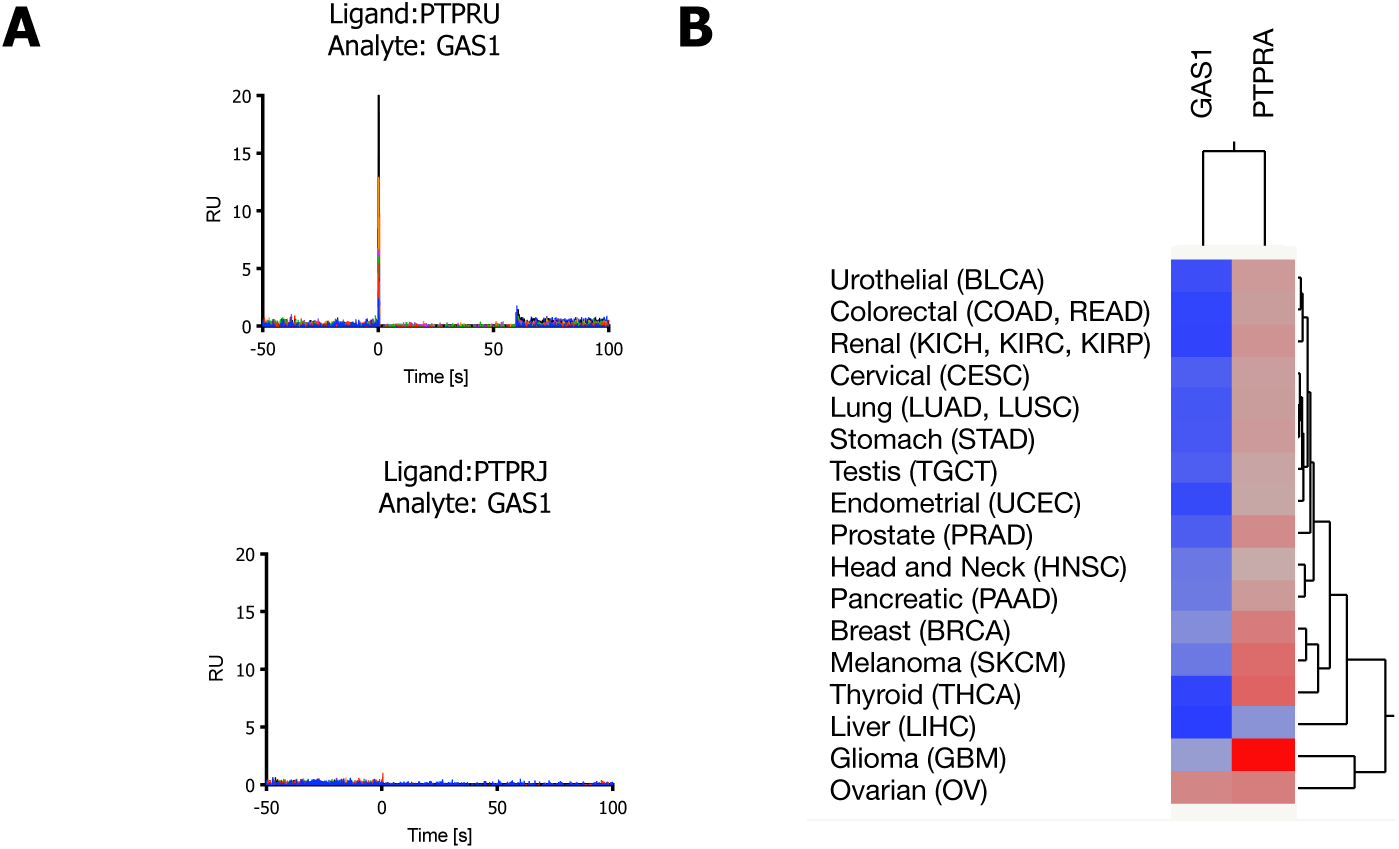
(A) SPR sensorgrams for testing binding of GAS1 to PTPRU or PRPRJ. (B) Unsupervised hierarchical clustering of normalized mRNA gene expression by tissue (normal vs. TCGA) was performed with Ward linkage and correlation distance were plotted as heatmaps using JMP.

**Figure S5.**
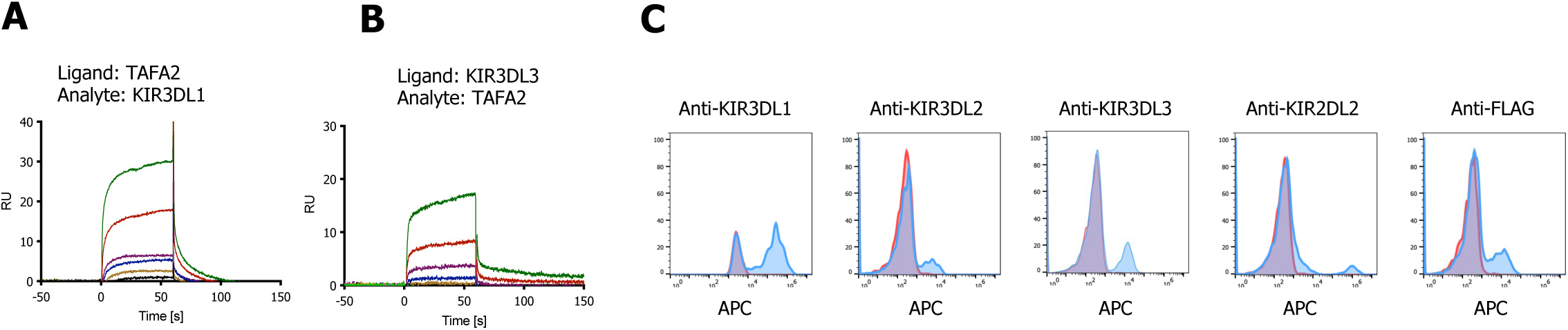
(A, B) SPR sensorgrams for binding of TAFA-2 to KIR3DL1 and KIR3DL3 ECDs respectively. (C) K562 cells lenti-virally transduced with KIR3DL1, KIR3DL2, KIR3DL3, KIR2DL2 or KIR2DL5A (FLAG-tagged) stained with the indicated APC-labelled antibodies (blue) compared to untransduced K562 parental cells (red).

**Figure S6.**
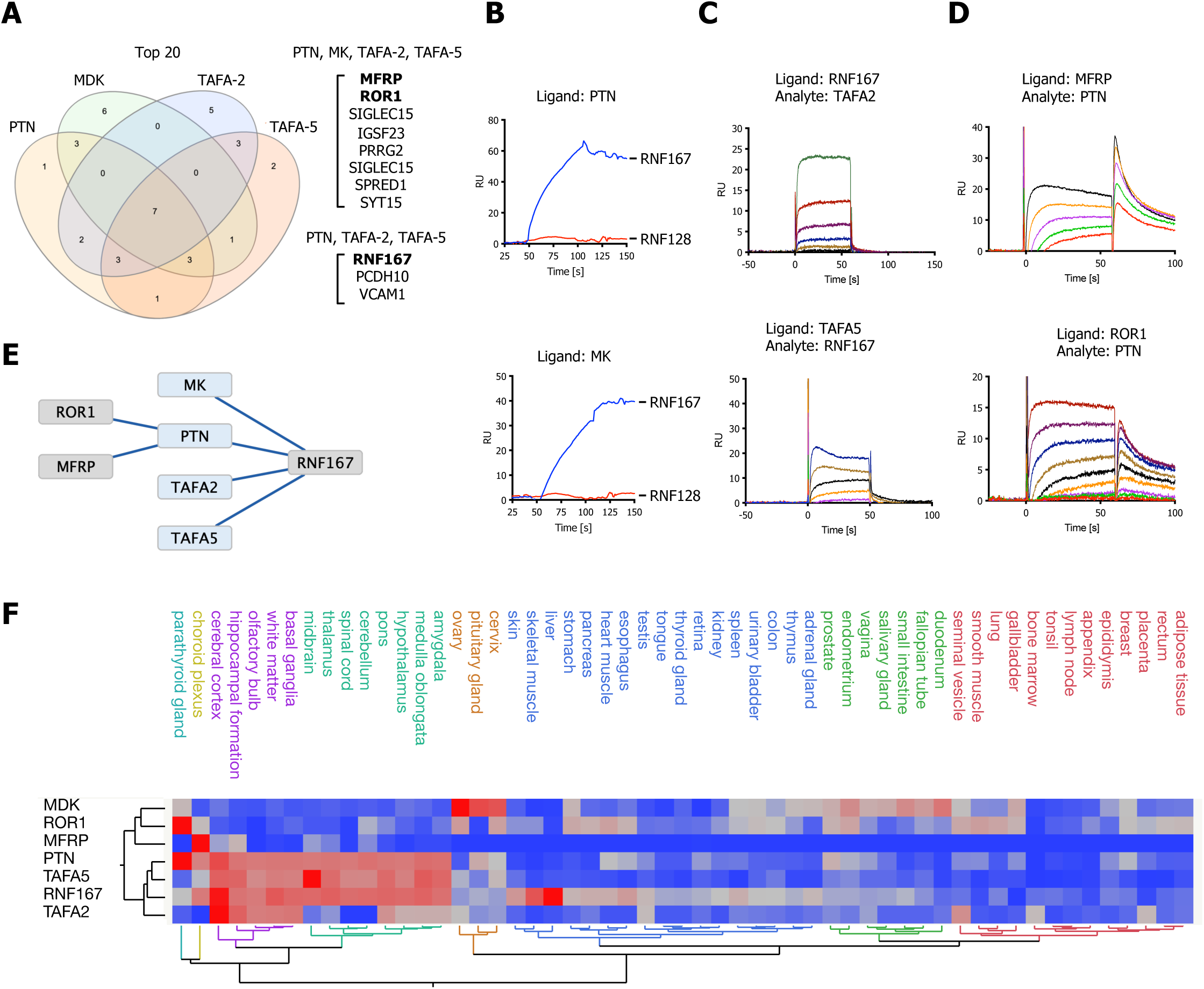
(A) Venn diagram depicting overlap of the top 20 ranking candidates for PTN, MK, TAFA-2 and TAFA-5 screens. Shared hits for PTN, MK, TAFA-2, TAFA-5 and shared candidates for PTN, TAFA-2 and TAFA-5 screens are listed. (B) MK or PTN ectodomains were captured on sensors (ligands) and analyzed for binding to RNF167 (RNF128 was used as a negative control). (C) SPR sensorgrams for human TAFA-2 and TAFA-5 binding to RNF167-ECD. (D) SPR sensorgrams for human PTN binding to MFRP-ECD and ROR1-ECD. (E) Visualization of the reported interactions in a node/edge network format. (F) Hierarchical 2-way clustering of mRNA expression data from normal tissue for PTN, MK, TAFA-2, TAFA-5, RNF167, MFRP and ROR1.

**Figure S7.**
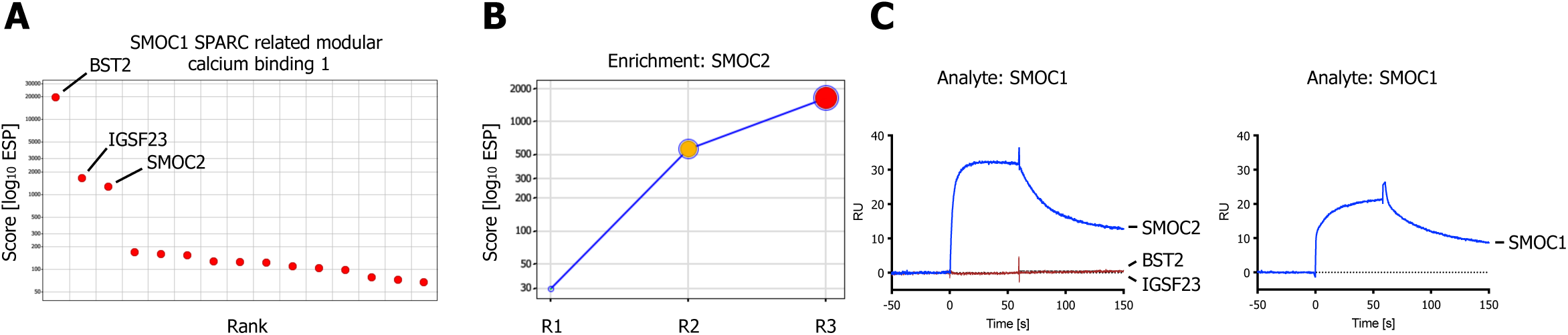
(A) SMOC1 ranked x/y scatter plot (ESP scores). (B) Trajectory plot of SMOC2 for all 3 consecutive rounds of selections in a x/y enrichment plot, size of the bubble represents the p-Value (-log10). (C) BST2, IGSF23 and SMOC2 ECDs were captured on sensors (ligands) and analyzed for binding to SMOC1 (analyte), SPR assay showing binding of SMOC1 (analyte) to SMOC2 or SMOC1, immobilized on a sensor chip (ligand).

**Figure S8.**
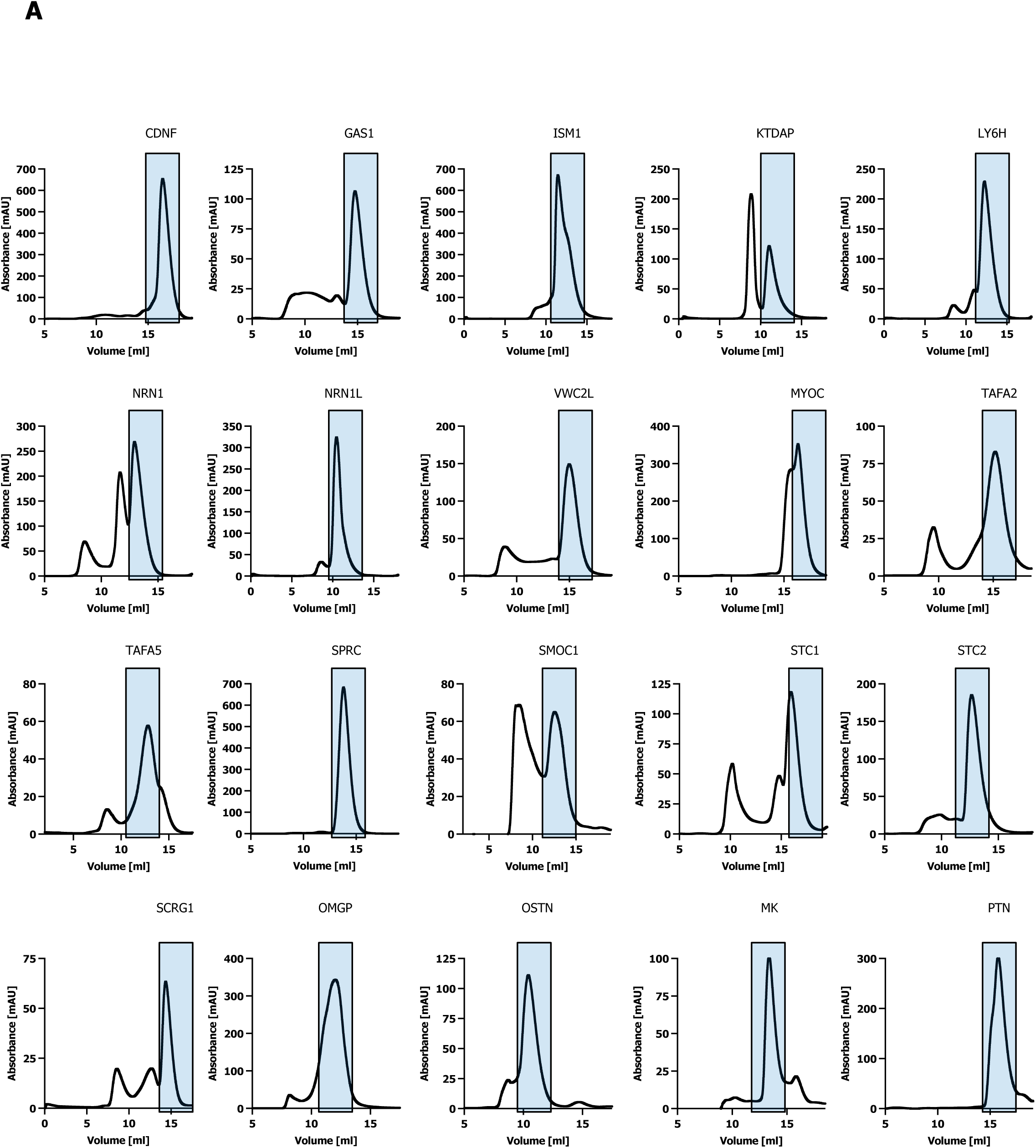
(A) Size-Exclusion Chromatography of secreted ligand bait proteins.

**Figure S9.**
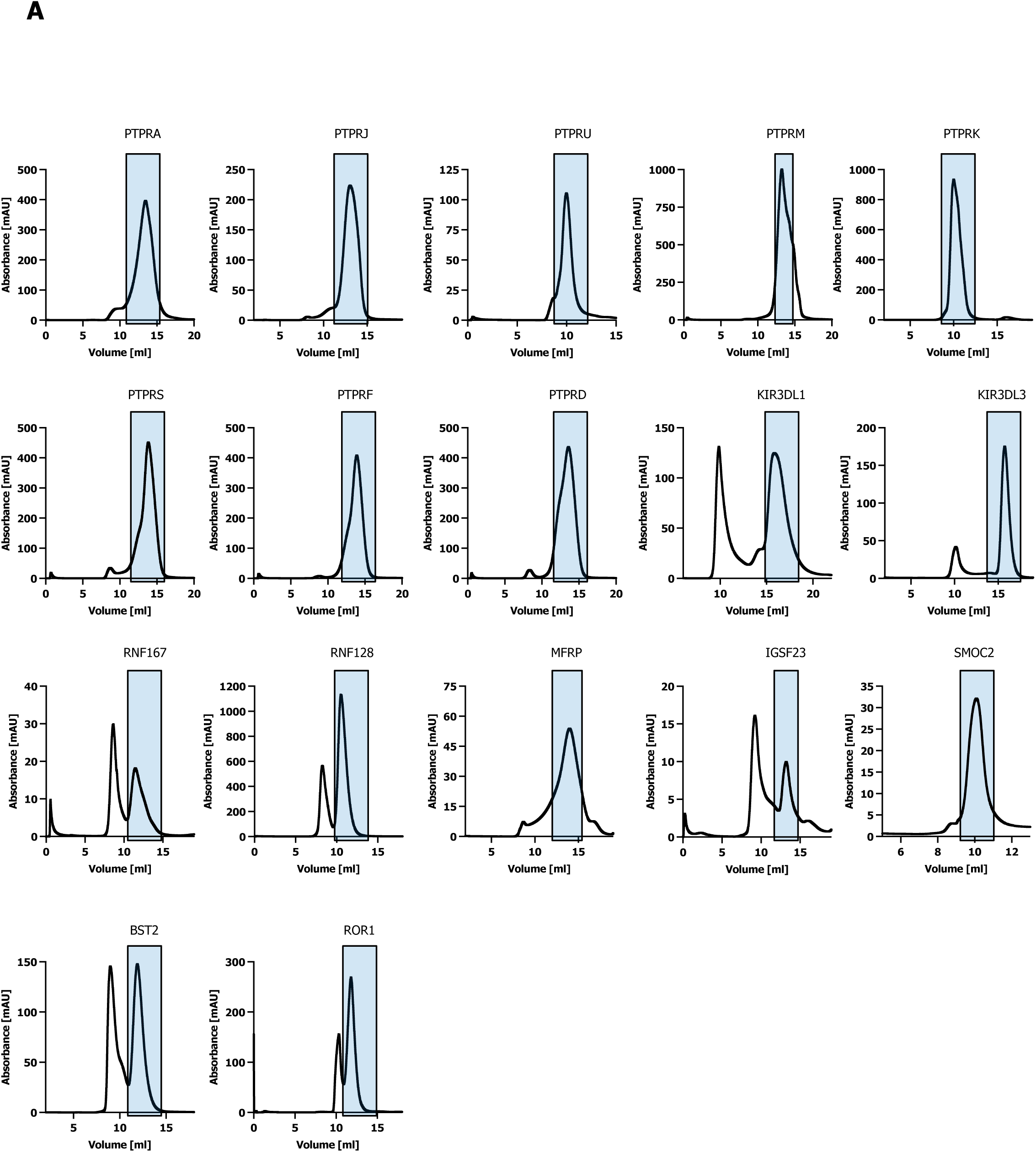
(A) Size-Exclusion Chromatography of proteins used for SPR.

## REFERENCES

1. Almagro Armenteros JJ, Tsirigos KD, Sønderby CK, Petersen TN, Winther O, Brunak S, von Heijne G, Nielsen H. 2019. SignalP 5.0 improves signal peptide predictions using deep neural networks. Nature Biotechnology. doi:10.1038/s41587-019-0036-z

2. Ataman B, Boulting GL, Harmin DA, Yang MG, Baker-Salisbury M, Yap EL, Malik AN, Mei K, Rubin AA, Spiegel I, Durresi E, Sharma N, Hu LS, Pletikos M, Griffith EC, Partlow JN, Stevens CR, Adli M, Chahrour M, Sestan N, Walsh CA, Berezovskii VK, Livingstone MS, Greenberg ME. 2016. Evolution of Osteocrin as an activity-regulated factor in the primate brain. Nature 539:242–247. doi:10.1038/NATURE20111

3. Braun P, Tasan M, Dreze M, Barrios-Rodiles M, Lemmens I, Yu H, Sahalie JM, Murray RR, Roncari L, de Smet AS, Venkatesan K, Rual JF, Vandenhaute J, Cusick ME, Pawson T, Hill DE, Tavernier J, Wrana JL, Roth FP, Vidal M. 2009. An experimentally derived confidence score for binary protein-protein interactions. Nat Methods 6:91–97. doi:10.1038/NMETH.1281

4. Brellier F, Ruggiero S, Zwolanek D, Martina E, Hess D, Brown-Luedi M, Hartmann U, Koch M, Merlo A, Lino M, Chiquet-Ehrismann R. 2011. SMOC1 is a tenascin-C interacting protein over-expressed in brain tumors. Matrix Biology 30:225–233. doi:10.1016/J.MATBIO.2011.02.001

5. Bushell KM, Söllner C, Schuster-Boeckler B, Bateman A, Wright GJ. 2008. Large-scale screening for novel low-affinity extracellular protein interactions. Genome Research. doi:10.1101/gr.7187808

6. Chong ZS, Ohnishi S, Yusa K, Wright GJ. 2018. Pooled extracellular receptor-ligand interaction screening using CRISPR activation. Genome Biology. doi:10.1186/s13059-018-1581-3

7. Clark HF, Gurney AL, Abaya E, Baker K, Baldwin D, Brush J, Chen J, Chow B, Chui C, Crowley C, Currell B, Deuel B, Dowd P, Eaton D, Foster J, Grimaldi C, Gu Q, Hass PE, Heldens S, Huang A, Kim HS, Klimowski L, Jin Y, Johnson S, Lee J, Lewis L, Liao D, Mark M, Robbie E, Sanchez C, Schoenfeld J, Seshagiri S, Simmons L, Singh J, Smith V, Stinson J, Vagts A, Vandlen R, Watanabe C, Wieand D, Woods K, Xie MH, Yansura D, Yi S, Yu G, Yuan J, Zhang M, Zhang Z, Goddard A, Wood WI, Godowski P. 2003. The secreted protein discovery initiative (SPDI), a large-scale effort to identify novel human secreted and transmembrane proteins: A bioinformatics assessment. Genome Research. doi:10.1101/gr.1293003

8. Cong L, Ran FA, Cox D, Lin S, Barretto R, Habib N, Hsu PD, Wu X, Jiang W, Marraffini LA, Zhang F. 2013. Multiplex genome engineering using CRISPR/Cas systems. Science (1979). doi:10.1126/science.1231143

9. Deans RM, Morgens DW, Ökesli A, Pillay S, Horlbeck MA, Kampmann M, Gilbert LA, Li A, Mateo R, Smith M, Glenn JS, Carette JE, Khosla C, Bassik MC. 2016. Parallel shRNA and CRISPR-Cas9 screens enable antiviral drug target identification. Nature Chemical Biology 12. doi:10.1038/nchembio.2050

10. Dobri AM, Dudău M, Enciu AM, Hinescu ME. 2021. CD36 in Alzheimer’s Disease: An Overview of Molecular Mechanisms and Therapeutic Targeting. Neuroscience 453:301–311. doi:10.1016/J.NEUROSCIENCE.2020.11.003

11. Doench JG. 2018. Am I ready for CRISPR? A user’s guide to genetic screens. Nat Rev Genet 19:67–80. doi:10.1038/NRG.2017.97

12. Endo M, Kamizaki K, Minami Y. 2022. The Ror-Family Receptors in Development, Tissue Regeneration and Age-Related Disease. Front Cell Dev Biol 10. doi:10.3389/FCELL.2022.891763

13. Ghilarducci K, Cabana VC, Desroches C, Chabi K, Bourgault S, Cappadocia L, Lussier MP. 2021. Functional interaction of ubiquitin ligase RNF167 with UBE2D1 and UBE2N promotes ubiquitination of AMPA receptor. FEBS Journal 288:4849–4868. doi:10.1111/febs.15796

14. Gilbert LA, Horlbeck MA, Adamson B, Villalta JE, Chen Y, Whitehead EH, Guimaraes C, Panning B, Ploegh HL, Bassik MC, Qi LS, Kampmann M, Weissman JS. 2014a. Genome-Scale CRISPR-Mediated Control of Gene Repression and Activation. Cell. doi:10.1016/j.cell.2014.09.029

15. Gilbert LA, Horlbeck MA, Adamson B, Villalta JE, Chen Y, Whitehead EH, Guimaraes C, Panning B, Ploegh HL, Bassik MC, Qi LS, Kampmann M, Weissman JS. 2014b. Genome-Scale CRISPR-Mediated Control of Gene Repression and Activation. Cell 159:647–661. doi:10.1016/J.CELL.2014.09.029

16. Grajchen E, Wouters E, van de Haterd B, Haidar M, Hardonnière K, Dierckx T, van Broeckhoven J, Erens C, Hendrix S, Kerdine-Römer S, Hendriks JJA, Bogie JFJ. 2020. CD36-mediated uptake of myelin debris by macrophages and microglia reduces neuroinflammation. J Neuroinflammation 17. doi:10.1186/S12974-020-01899-X

17. Green J, Nusse R, van Amerongen R. 2014. The role of Ryk and Ror receptor tyrosine kinases in Wnt signal transduction. Cold Spring Harbor Perspectives in Biology 6.

18. Havugimana PC, Hart GT, Nepusz T, Yang H, Turinsky AL, Li Z, Wang PI, Boutz DR, Fong V, Phanse S, Babu M, Craig SA, Hu P, Wan C, Vlasblom J, Dar VUN, Bezginov A, Clark GW, Wu GC, Wodak SJ, Tillier ERM, Paccanaro A, Marcotte EM, Emili A. 2012. A census of human soluble protein complexes. Cell 150:1068–1081. doi:10.1016/J.CELL.2012.08.011

19. Hay IM, Fearnley GW, Rios P, Köhn M, Sharpe HJ, Deane JE. 2020. The receptor PTPRU is a redox sensitive pseudophosphatase. Nature Communications 2020 11:1 11:1–13. doi:10.1038/s41467-020-17076-w

20. Heberle H, Meirelles VG, da Silva FR, Telles GP, Minghim R. 2015. InteractiVenn: A web-based tool for the analysis of sets through Venn diagrams. BMC Bioinformatics 16:1–7. doi:10.1186/S12859-015-0611-3/FIGURES/4

21. Honig B, Shapiro L. 2020. Adhesion Protein Structure, Molecular Affinities, and Principles of Cell-Cell Recognition. Cell 181:520–535. doi:10.1016/J.CELL.2020.04.010

22. Huang S, Zheng C, Xie G, Song Z, Wang P, Bai Y, Chen D, Zhang Y, Lv P, Liang W, She S, Li Q, Liu Z, Wang Yun, Xing GG, Wang Ying. 2021. FAM19A5/TAFA5, a novel neurokine, plays a crucial role in depressive-like and spatial memory-related behaviors in mice. Mol Psychiatry 26:2363–2379. doi:10.1038/S41380-020-0720-X

23. Huttlin EL, Bruckner RJ, Paulo JA, Cannon JR, Ting L, Baltier K, Colby G, Gebreab F, Gygi MP, Parzen H, Szpyt J, Tam S, Zarraga G, Pontano-Vaites L, Swarup S, White AE, Schweppe DK, Rad R, Erickson BK, Obar RA, Guruharsha KG, Li K, Artavanis-Tsakonas S, Gygi SP, Wade Harper J. 2017. Architecture of the human interactome defines protein communities and disease networks. Nature. doi:10.1038/nature22366

24. Huttlin EL, Ting L, Bruckner RJ, Gebreab F, Gygi MP, Szpyt J, Tam S, Zarraga G, Colby G, Baltier K, Dong R, Guarani V, Vaites LP, Ordureau A, Rad R, Erickson BK, Wühr M, Chick J, Zhai B, Kolippakkam D, Mintseris J, Obar RA, Harris T, Artavanis-Tsakonas S, Sowa ME, de Camilli P, Paulo JA, Harper JW, Gygi SP. 2015. The BioPlex Network: A Systematic Exploration of the Human Interactome. Cell 162:425–440.

25. Jinek M, East A, Cheng A, Lin S, Ma E, Doudna J. 2013. RNA-programmed genome editing in human cells. Elife. doi:10.7554/eLife.00471

26. Julien SG, Dubé N, Hardy S, Tremblay ML. 2010. Inside the human cancer tyrosine phosphatome. Nature Reviews Cancer 2011 11:1 11:35–49. doi:10.1038/nrc2980

27. Kampmann M. 2018. CRISPRi and CRISPRa Screens in Mammalian Cells for Precision Biology and Medicine. ACS Chemical Biology 13:406–416. doi:10.1021/acschembio.7b00657

28. Karlsson M, Zhang C, Méar L, Zhong W, Digre A, Katona B, Sjöstedt E, Butler L, Odeberg J, Dusart P, Edfors F, Oksvold P, von Feilitzen K, Zwahlen M, Arif M, Altay O, Li X, Ozcan M, Mardonoglu A, Fagerberg L, Mulder J, Luo Y, Ponten F, Uhlén M, Lindskog C. 2021. A single–cell type transcriptomics map of human tissues. Science Advances 7. doi:10.1126/SCIADV.ABH2169/SUPPL_FILE/SCIADV.ABH2169_SM.PDF

29. Katoh M. 2001. Molecular cloning and characterization of MFRP, a novel gene encoding a membrane-type Frizzled-related protein. Biochem Biophys Res Commun 282:116–123. doi:10.1006/BBRC.2001.4551

30. Krogan NJ, Cagney G, Yu H, Zhong G, Guo X, Ignatchenko A, Li J, Pu S, Datta N, Tikuisis AP, Punna T, Peregrín-Alvarez JM, Shales M, Zhang X, Davey M, Robinson MD, Paccanaro A, Bray JE, Sheung A, Beattie B, Richards DP, Canadien V, Lalev A, Mena F, Wong P, Starostine A, Canete MM, Vlasblom J, Wu S, Orsi C, Collins SR, Chandran S, Haw R, Rilstone JJ, Gandi K, Thompson NJ, Musso G, St Onge P, Ghanny S, Lam MHY, Butland G, Altaf-Ul AM, Kanaya S, Shilatifard A, O’Shea E, Weissman JS, Ingles CJ, Hughes TR, Parkinson J, Gerstein M, Wodak SJ, Emili A, Greenblatt JF. 2006. Global landscape of protein complexes in the yeast Saccharomyces cerevisiae. Nature 440:637–643. doi:10.1038/NATURE04670

31. Li H, Watson A, Olechwier A, Anaya M, Sorooshyari SK, Harnett DP, Lee HK (Peter), Vielmetter J, Fares MA, Garcia KC, Özkan E, Labrador JP, Zinn K. 2017. Deconstruction of the beaten path-sidestep interaction network provides insights into neuromuscular system development. Elife 6. doi:10.7554/ELIFE.28111

32. Li Y, Mariuzza RA. 2014. Structural Basis for Recognition of Cellular and Viral Ligands by NK Cell Receptors. Frontiers in Immunology 5. doi:10.3389/FIMMU.2014.00123

33. Long F, Shi H, Li P, Guo S, Ma Y, Wei S, Li Y, Gao F, Gao S, Wang M, Duan R, Wang X, Yang K, Sun W, Li X, Li J, Liu Q. 2021. A SMOC2 variant inhibits BMP signaling by competitively binding to BMPR1B and causes growth plate defects. Bone 142:115686. doi:10.1016/J.BONE.2020.115686

34. Lussier MP, Herring BE, Nasu-Nishimura Y, Neutzner A, Karbowski M, Youle RJ, Nicoll RA, Roche KW. 2012. Ubiquitin ligase RNF167 regulates AMPA receptor-mediated synaptic transmission. Proc Natl Acad Sci U S A 109:19426–19431. doi:10.1073/PNAS.1217477109/-/DCSUPPLEMENTAL

35. Mali P, Yang L, Esvelt KM, Aach J, Guell M, DiCarlo JE, Norville JE, Church GM. 2013. RNA-guided human genome engineering via Cas9. Science (1979). doi:10.1126/science.1232033

36. Martinez-Martin N. 2017. Technologies for Proteome-Wide Discovery of Extracellular Host-Pathogen Interactions. J Immunol Res 2017:2197615–2197618.

37. Martinez-Martin N, Verschueren E, Husain B. 2019. The Immunoglobulin Superfamily Receptome Defines Cancer-Relevant Networks Associated with Response to Immunotherapy. Annals of Oncology. doi:10.1093/annonc/mdz451.004

38. Minami Y, Oishi I, Endo M, Nishita M. 2010. Ror-family receptor tyrosine kinases in noncanonical Wnt signaling: Their implications in developmental morphogenesis and human diseases. Developmental Dynamics 239:1–15. doi:10.1002/dvdy.21991

39. Moffatt P, Thomas G, Sellin K, Bessette MC, Lafrenière F, Akhouayri O, St-Arnaud R, Lanctôt C. 2007. Osteocrin is a specific ligand of the natriuretic Peptide clearance receptor that modulates bone growth. J Biol Chem 282:36454–36462. doi:10.1074/JBC.M708596200

40. Morgens DW, Deans RM, Li A, Bassik MC. 2016. Systematic comparison of CRISPR/Cas9 and RNAi screens for essential genes. Nature Biotechnology. doi:10.1038/nbt.3567

41. Nakamura N. 2011. The Role of the Transmembrane RING Finger Proteins in Cellular and Organelle Function. Membranes 2011, Vol 1, Pages 354–393 1:354–393. doi:10.3390/MEMBRANES1040354

42. Özkan E, Carrillo RA, Eastman CL, Weiszmann R, Waghray D, Johnson KG, Zinn K, Celniker SE, Garcia KC. 2013a. An extracellular interactome of immunoglobulin and LRR proteins reveals receptor-ligand networks. Cell 154:228. doi:10.1016/J.CELL.2013.06.006

43. Özkan E, Carrillo RA, Eastman CL, Weiszmann R, Waghray D, Johnson KG, Zinn K, Celniker SE, Garcia KC. 2013b. XAn extracellular interactome of immunoglobulin and LRR proteins reveals receptor-ligand networks. Cell. doi:10.1016/j.cell.2013.06.006

44. Özkan E, Özkan E, Carrillo RA, Eastman CL, Weiszmann R, Waghray D, Johnson KG, Zinn K, Celniker SE, Garcia KC, Garcia KC. 2013c. An extracellular interactome of immunoglobulin and LRR proteins reveals receptor-ligand networks. Cell 154:228–239.

45. Pende D, Falco M, Vitale M, Cantoni C, Vitale C, Munari E, Bertaina A, Moretta F, del Zotto G, Pietra G, Mingari MC, Locatelli F, Moretta L. 2019. Killer Ig-Like Receptors (KIRs): Their Role in NK Cell Modulation and Developments Leading to Their Clinical Exploitation. Front Immunol 10. doi:10.3389/FIMMU.2019.01179

46. Pierleoni A, Martelli P, Casadio R. 2008. PredGPI: A GPI-anchor predictor. BMC Bioinformatics. doi:10.1186/1471-2105-9-392

47. Ranaivoson FM, Turk LS, Ozgul S, Kakehi S, von Daake S, Lopez N, Trobiani L, de Jaco A, Denissova N, Demeler B, Özkan E, Montelione GT, Comoletti D. 2019. A Proteomic Screen of Neuronal Cell-Surface Molecules Reveals IgLONs as Structurally Conserved Interaction Modules at the Synapse. Structure 27:893–906.e9. doi:10.1016/J.STR.2019.03.004

48. Sarver DC, Lei X, Wong GW. 2021. FAM19A (TAFA): An Emerging Family of Neurokines with Diverse Functions in the Central and Peripheral Nervous System. ACS Chem Neurosci 12:945–958. doi:10.1021/ACSCHEMNEURO.0C00757

49. Silverstein RL, Febbraio M. 2009. CD36, a Scavenger Receptor Involved in Immunity, Metabolism, Angiogenesis, and Behavior. Sci Signal 2:re3. doi:10.1126/SCISIGNAL.272RE3

50. Sivori S, Vacca P, del Zotto G, Munari E, Mingari MC, Moretta L. 2019. Human NK cells: surface receptors, inhibitory checkpoints, and translational applications. Cellular and Molecular Immunology 16:430–441. doi:10.1038/s41423-019-0206-4

51. Söllner C, Wright GJ. 2009. A cell surface interaction network of neural leucine-rich repeat receptors. Genome Biol 10:R99–R99.

52. Stastna M, Van Eyk JE. 2012. Secreted proteins as a fundamental source for biomarker discovery. Proteomics. doi:10.1002/pmic.201100346

53. Subbotina E, Sierra A, Zhu Z, Gao Z, Koganti SRK, Reyes S, Stepniak E, Walsh SA, Acevedo MR, Perez-Terzic CM, Hodgson-Zingman DM, Zingman L v. 2015. Musclin is an activity-stimulated myokine that enhances physical endurance. Proc Natl Acad Sci U S A 112:16042–16047. doi:10.1073/PNAS.1514250112

54. Sundin OH, Dharmaraj S, Bhutto IA, Hasegawa T, McLeod DS, Merges CA, Silval ED, Maumenee IH, Lutty GA. 2008. Developmental basis of nanophthalmos: MFRP Is required for both prenatal ocular growth and postnatal emmetropization. Ophthalmic Genet 29:1–9. doi:10.1080/13816810701651241

55. Tanenbaum ME, Gilbert LA, Qi LS, Weissman JS, Vale RD. 2014a. A Protein-Tagging System for Signal Amplification in Gene Expression and Fluorescence Imaging. Cell 159:635–646.

56. Tanenbaum ME, Gilbert LA, Qi LS, Weissman JS, Vale RD. 2014b. A Protein-Tagging System for Signal Amplification in Gene Expression and Fluorescence Imaging. Cell 159:635–646.

57. Taouji S, Dahan S, Bosse R, Chevet E. 2009. Current Screens Based on the AlphaScreen Technology for Deciphering Cell Signalling Pathways. Curr Genomics 10:93–101. doi:10.2174/138920209787847041

58. Thul PJ, Akesson L, Wiking M, Mahdessian D, Geladaki A, Ait Blal H, Alm T, Asplund A, Björk L, Breckels LM, Bäckström A, Danielsson F, Fagerberg L, Fall J, Gatto L, Gnann C, Hober S, Hjelmare M, Johansson F, Lee S, Lindskog C, Mulder J, Mulvey CM, Nilsson P, Oksvold P, Rockberg J, Schutten R, Schwenk JM, Sivertsson A, Sjöstedt E, Skogs M, Stadler C, Sullivan DP, Tegel H, Winsnes C, Zhang C, Zwahlen M, Mardinoglu A, Pontén F, Von Feilitzen K, Lilley KS, Uhlén M, Lundberg E. 2017. A subcellular map of the human proteome. Science (1979). doi:10.1126/science.aal3321

59. Tom Tang Y, Emtage P, Funk WD, Hu T, Arterburn M, Park EEJ, Rupp F. 2004. TAFA: a novel secreted family with conserved cysteine residues and restricted expression in the brain. Genomics 83:727–734. doi:10.1016/J.YGENO.2003.10.006

60. Tonks NK. 2006. Protein tyrosine phosphatases: from genes, to function, to disease. Nature Reviews Molecular Cell Biology 7:833–846.

61. Uhlén M, Karlsson MJ, Hober A, Svensson AS, Scheffel J, Kotol D, Zhong W, Tebani A, Strandberg L, Edfors F, Sjöstedt E, Mulder J, Mardinoglu A, Berling A, Ekblad S, Dannemeyer M, Kanje S, Rockberg J, Lundqvist M, Malm M, Volk AL, Nilsson P, Månberg A, Dodig-Crnkovic T, Pin E, Zwahlen M, Oksvold P, von Feilitzen K, Häussler RS, Hong MG, Lindskog C, Ponten F, Katona B, Vuu J, Lindström E, Nielsen J, Robinson J, Ayoglu B, Mahdessian D, Sullivan D, Thul P, Danielsson F, Stadler C, Lundberg E, Bergström G, Gummesson A, Voldborg BG, Tegel H, Hober S, Forsström B, Schwenk JM, Fagerberg L, Sivertsson Å. 2019. The human secretome. Sci Signal 12. doi:10.1126/SCISIGNAL.AAZ0274

62. Verschueren E, Husain B, Yuen K, Sun Y, Paduchuri S, Senbabaoglu Y, Lehoux I, Arena TA, Wilson B, Lianoglou S, Bakalarski C, Franke Y, Chan P, Wong AW, Gonzalez LC, Mariathasan S, Turley SJ, Lill JR, Martinez-Martin N. 2020. The Immunoglobulin Superfamily Receptome Defines Cancer-Relevant Networks Associated with Clinical Outcome. Cell 182:329–344.e19. doi:10.1016/J.CELL.2020.06.007

63. Wojtowicz WM, Vielmetter J, Fernandes RA, Siepe DH, Eastman CL, Chisholm GB, Cox S, Klock H, Anderson PW, Rue SM, Miller JJ, Glaser SM, Bragstad ML, Vance J, Lam AW, Lesley SA, Zinn K, Garcia KC. 2020. A Human IgSF Cell-Surface Interactome Reveals a Complex Network of Protein-Protein Interactions. Cell. doi:10.1016/j.cell.2020.07.025

64. Yan J, Jia H, Ma Z, Ye H, Zhou M, Su L, Liu J, Guo AY. 2014. The evolutionary analysis reveals domain fusion of proteins with Frizzled-like CRD domain. Gene 533:229–239. doi:10.1016/J.GENE.2013.09.083

